# Impairing one sensory modality enhances another by reprogramming peptidergic circuits in *Caenorhabditis elegans*

**DOI:** 10.1101/2021.03.11.435052

**Authors:** Giulio Valperga, Mario de Bono

## Abstract

Animals that lose one sensory modality often show augmented responses to other sensory inputs. The mechanisms underpinning this cross-modal plasticity are poorly understood. To probe these mechanisms, we perform a forward genetic screen for mutants with enhanced O_2_ perception in *C. elegans*. Multiple mutants exhibiting increased responsiveness to O_2_ concomitantly show defects in other sensory responses. One mutant, *qui-1*, defective in a conserved NACHT/WD40 protein, abolishes pheromone-evoked Ca^2+^ responses in the ADL chemosensory neurons. We find that ADL’s responsiveness to pre-synaptic input from O_2_- sensing neurons is heightened in *qui-1* and other sensory defective mutants resulting in an enhanced neurosecretion. Expressing *qui-1* selectively in ADL rescues both the *qui-1* ADL neurosecretory phenotype and enhanced escape from 21% O_2_. Profiling of ADL neurons indicates its acquired O_2_-evoked neurosecretion is the result of a transcriptional reprogramming that up-regulates neuropeptide signalling. We show that the conserved neuropeptide receptor NPR-22 is necessary and sufficient in ADL to enhance its neurosecretion levels. Sensory loss can thus confer cross-modal plasticity by re-wiring peptidergic circuits.

## Introduction

Animals that lose a sensory modality can adapt to the loss by increasing their sensitivity to other sensory inputs. This change can involve re-purposing neurons or brain areas that normally mediate responses to the lost modality such that they process other sensory inputs. For example, in blind people the absence of visual stimulation promotes rewiring of inputs into primary visual cortex (V1), so that it becomes responsive to tactile stimuli, a characteristic absent in sighted individuals (Büchel et al., 1998; Dietrich et al., 2013; Sadato et al., 1996; Wanet-Defalque et al., 1988). The molecular mechanisms enabling such re-purposing of neural circuits are incompletely understood, but at some level are thought to reflect opportunities for rewiring. More importantly, because of the heterogenic and complex aetiology of blindness, it is unclear how genetic lesions that result in sensory loss promote cross modal plasticity (Merabet and Pascual-Leone, 2009). Animals can execute innate behaviors without a need for prior learning. However, experience and context can modulate innate behaviors, with circuits coordinating innate responses integrating information from modulating sensory pathways. These connections can link circuits that mediate responses to distinct sensory cues, thus providing opportunities for plastic changes to re-route sensory information if one sensory pathway is damaged (Fine and Park, 2016). *C. elegans* provides a favourable model to study cross-modal interactions in neural circuits, and how these connections may be altered by neural plasticity. This is possible thanks to a careful reconstruction of its complete wiring diagram of chemical and electrical synapses (Cook et al., 2019; Jarrell et al., 2012; White et al., 1986). These studies have emphasized the anatomical stereotypy of the *C. elegans* nervous system which contrasts with extensive experience-dependent plasticity at the behavioral level (Fenk and de Bono, 2017; Pocock and Hobert, 2010; Saeki et al., 2001; Zhang et al., 2005).

A salient environmental cue for *C. elegans* is oxygen (O_2_) levels. Instantaneous as well as prior O_2_experience can reconfigure the value of sensory cues for this animal. For example, animals acclimated to 21% O_2_are attracted to pheromones that repel animals acclimated to 7% O_2_ (Fenk and de Bono, 2017). The wiring diagram, coupled with Ca^2+^ imaging, provide tantalizing hints about the basis of cross-modal plasticity associated with changes in O_2_ levels. One of the main O_2_-sensing neurons, URX, forms a spoke in a large hub-and-spoke circuit centred on the RMG interneurons (Macosko et al., 2009). Several sensory neurons, including pheromone receptors called ASK and ADL that respectively mediate attraction and repulsion from pheromones, form additional spokes in the circuit (Jang et al., 2012; Macosko et al., 2009). The URX O_2_ sensors show persistent higher activity at 21% O_2_compared to 7% O_2_, and tonically transmit this activity to the RMG hub interneurons (Busch et al., 2012). These O_2_-evoked changes in URX and RMG somehow reprogram the pheromone response properties of ASK and ADL (Fenk and de Bono, 2017). Reciprocally, altering sensory transduction in the ASK or ADL neurons influences how *C. elegans* responds to O_2_ stimuli (de Bono et al., 2002; Laurent et al., 2015; Macosko et al., 2009). However, the molecular underpinnings of how cross-modal changes are coordinated across the hub-and-spoke circuit as different elements of the circuit become more or less active are unclear.

Here, we employ forward genetics to identify mechanisms that promote cross-modal plasticity across the RMG hub-and-spoke circuit in response to genetic lesions. Since the circuit integrates several sensory modalities, we use genetic backgrounds that reduce signalling across the hub-and-spoke circuit and suppress *C. elegans* arousal in response to 21% O_2_. Mutations that promote cross-modal plasticity are likely to restore its O_2_-responses. We identify several sensory defective mutants that increase ADL’s ability to transmit information from its pre-synaptic O_2_ sensors, including URX and RMG. Specifically, the mutants show increased O_2_-evoked secretion of neuropeptides from ADL in response to oxygen inputs. Using RNAseq we profile ADL neurons in wild type and one enhancer mutant, *qui-1*, and discover rewiring of peptidergic circuits. Reprogramming of ADL’s peptidergic proprieties increase expression of the neuropeptide receptor NPR-22, which is necessary and sufficient to increase neurosecretion from ADL. In summary, our data suggest that loss of sensory reception in the ADL pheromone sensors increases ADL’s responsiveness to input from the O_2_-circuit by reprogramming its peptidergic properties. Rewiring peptidergic signalling across circuits may be an unappreciated mechanism by which loss of one sensory modality enhances responsiveness to another.

## Results

### A genetic screen for enhancers of *C. elegans* aggregation behavior

Wild isolates of *C. elegans* avoid and escape 21% O_2_ (de Bono and Bargmann, 1998). On a bacterial lawn these animals move rapidly and continuously while seeking lower O_2_ concentrations such as areas of thick bacterial growth. A hub-and-spoke network that integrates multiple sensory cues coordinates this escape behavior (Fenk and de Bono, 2017; Laurent et al., 2015; Macosko et al., 2009). The standard *C. elegans* lab strain N2 is not aroused by 21% O_2_, and does not accumulate on thick bacteria, due to a gain-of-function mutation in the neuropeptide receptor NPR-1 (de Bono and Bargmann, 1998). NPR-1 inhibits these behaviors by acting in the RMG interneurons to reduce the outputs of the hub-and-spoke circuit (Macosko et al., 2009). Since enhanced responsiveness to one sensory modality is a characteristic feature of cross-modal plasticity, we reasoned mutants with a restored O_2_- escape behaviour should allow us to isolate animals showing cross-modal plasticity.

To identify such mutants, we mutagenized N2 animals and selected for individuals that preferentially accumulated on a patch of thick bacteria (OP50) placed in the middle of a thin lawn (see Methods) (Figure 1A – Step 1). Besides N2, we mutagenized animals defective in *flp-21*, which encodes a neuropeptide ligand of the NPR-1 receptor. Disrupting *flp-21* weakly enhances O_2_ escape behavior (Laurent et al., 2015; Rogers et al., 2003), providing a sensitized background. We isolated 22 mutants from the N2 parental strain and 17 mutants from the *flp-21* parental strain that preferentially accumulated on the thick food patch.

**Figure 1.**
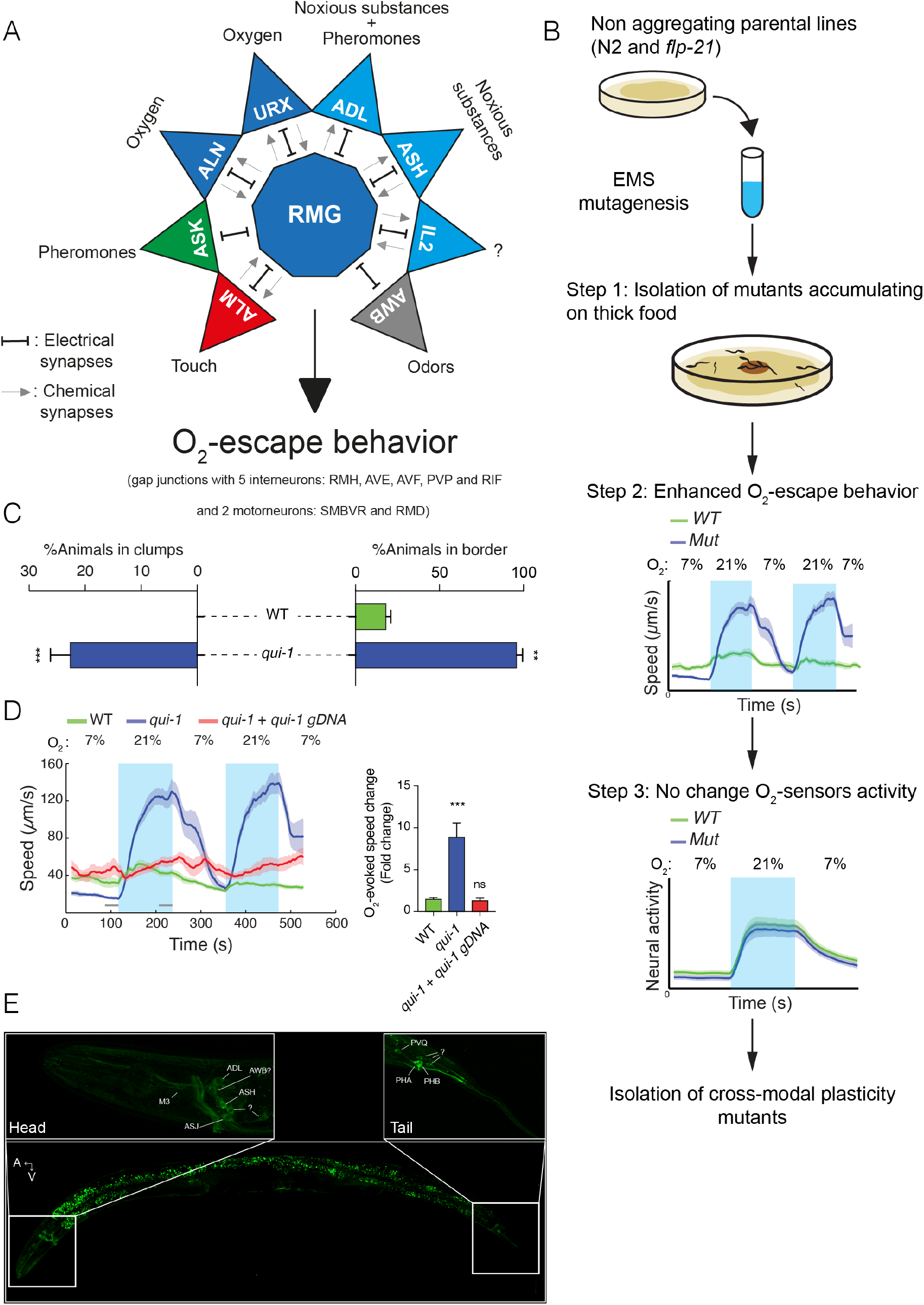
A genetic screen for mutations that induce cross-modal plasticity.(**A**) The hub and spoke circuit associated with the URX O_2_ sensors and O_2_ escape behavior. Updated version according to (Cook et al., 2019). (**B**) Schematic of the genetic screen. We selected mutants that preferentially accumulate on thicker bacteria, a behaviour that depends on oxygen responses (Step 1), screened these for increased O_2_-escape behavior (Step 2), and then identified strains with overtly normal O_2_-evoked Ca^2+^ responses in the URX O_2_-sensors and RMG interneurons (Step 3). (**C**) Bar graphs quantifying aggregation and bordering behavior. N=4–6 assays. (**D**) A wild type copy of *qui-1* rescues the O_2_-escape phenotype of *qui-1(db104)* mutants. Left: Line shows average speed, shading shows standard error of the mean (SEM) and grey bars show 30s time intervals used to average the animal’s speed. Right: Bar graph shows the fold change in average speed at 21% O_2_ compared to 7% O_2_ N= 6–9 assays. (**E**) QUI-1 expression and localization in an *mNeonGreen::qui-1* translational fusion knock-in strain. Fluorescent neurons include ADL, ASH (Head) and PVQ, PHB and PHA (Tail), and potentially M3, AWB and ASJ based on position and morphology. Also visible is yellow gut autofluorescence. Statistics:**=p value ≤0.01, ***=p value ≤0.001, ns=not significant, Mann-Whitney U test. Comparisons are with WT.

To confirm our mutants also showed an elevated O_2_-escape behavior, we further selected from our collection 6 mutants that displayed a larger O_2_-escape behavior than wild type (Figure 1B – Step 2). To identify the genetic defects causing increased O_2_-escape behavior in these mutants we used a Deep Sequence Mapping strategy (See Methods and Zuryn et al., 2010). A list of *de novo* high impact mutations highlighted only one likely loss-of-function mutation, a premature stop-codon (Q966Stop), within the *qui-1* gene (Figure S1A). Previous work suggested *qui-1* mutants lay eggs where bacteria are thickest (Neal et al., 2016). *qui-1(db104)* mutants isolated in our screen displayed both aggregation and O_2_-escape behavior (Figure 1C-1D). We next compared the O_2_-escape behavior of the *db104* mutant with a strain carrying a deletion allele, *qui-1*(*ok3571*) (Figure S1A). These strains showed indistinguishable responses (Figure S1B), confirming *qui-1* (*db104*) is a loss-of-function mutation that leads to a strong O_2_-escape behavior. To confirm this further, we showed that a wild type *qui-1* transgene completely rescued *qui-1* (*db104*) O_2_-escape phenotype (Figure 1D).

In the hub-and-spoke circuit, the URX O_2_ sensors are tonically activated by 21% O_2_ and in turn tonically activate the RMG hub interneurons (Busch et al., 2012). Optogenetic experiments show that increasing URX or RMG activity is sufficient to stimulate rapid movement (Busch et al., 2012). To probe the *qui-1* phenotype, we imaged O_2_-evoked Ca^2+^ responses in URX and RMG neurons in *qui-1* mutants (Figure 1B – Step 3). *qui-1* Ca^2+^ responses did not differ from those of wild type controls (Figure S1C and S1D), suggesting that its augmented O_2_ escape behavior does not reflect a simple increase in URX or RMG activity.

### The NACHT/WD40 protein QUI-1 acts in the ASH and ADL spoke neurons to inhibit O_2_-escape behavior

Previous work (Hilliard et al., 2004) and homology searches suggest QUI-1 is an ortholog of NWD1 (Nacht and WD40 repeat domain containing 1), a conserved protein of poorly understood function (Figure S2A and S2B). The *C. elegans* genome also encodes a paralog of QUI-1, T05C3.2, most similar to mammalian NWD2 (Figure 2SB). These proteins combine a NACHT domain with multiple WD40 domains and have homologs across phylogeny (Figure S2B and S2C). WD40 domains mediate protein-protein or protein-DNA interactions. NACHT domains are present in proteins involved in programmed cell death and transcription of the Major Histocompatibility Complex (MHC), and include an NTPase domain, which is proposed to regulate signalling from these proteins. Most of the Walker A motif (Motif 1 - P loop) in the NTPase domain, which binds nucleotides, is conserved in QUI-1 (Figure S2C), suggesting the NACHT domain is functional.

Previous work suggests *qui-1* is expressed in a small subset of sensory and interneurons (Hilliard et al., 2004). To confirm *qui-1* expression pattern, we used CRISPR/Cas9 genome editing to insert DNA encoding the mNeonGreen fluorescent protein in frame with the N terminus of QUI-1. Fluorescence from the mNeonGreen::QUI-1 fusion protein was confined to head and tail neurons, and we confirmed expression in ASH, ADL, PHB and PVQ as previously reported (Figure 1E and Hilliard et al., 2004). We observed expression in five additional neurons close to the nerve ring, including possibly M3, AWB and ASJ, and three neurons close to the tail (Figure 1E). The mNeonGreen::QUI-1 fusion protein appears to be largely cytosolic and excluded from the nucleus, consistent with previous reports (Figure 1D and Neal et al., 2016).

Two of *qui-1*-expressing neurons, ASH and ADL form part of the RMG hub-and-spoke circuit (Macosko et al., 2009). ASH and ADL have previously been shown to promote aggregation and escape from 21% O_2_ (de Bono et al., 2002), although they are probably not primary O_2_ sensors. ASH and ADL are nociceptors that mediate *C. elegans* avoidance of a variety of chemical and non-chemical stimuli (Hilliard et al., 2005; Jang et al., 2012), e.g. Cu^2+^ (ASH) and pheromones (ADL). We used cell-specific rescue of *qui-1* mutants to ask if QUI-1 acts in ASH and/or ADL neurons to inhibit O_2_-evoked escape behavior. Expressing *qui-1* selectively in ASH neurons reduced the O_2_-escape response of *qui-1* mutants compared to controls (Figure 2A). The rescue was not complete: transgenic animals retained a significant O_2_-response compared to wild type controls (Figure 2A). Expressing *qui-1* only in ADL also significantly reduced the O_2_-evoked escape behavior of *qui-1* mutants (Figure 2B), but as with targeted expression in ASH, rescue was incomplete and transgenic animals responded significantly more to a 21% O_2_ stimulus than wild type controls (Figure 2B). Expressing QUI-1 in both ASH and ADL neurons did not show an additive rescue effect (Figure 2C), consistent with ablation studies suggesting these neurons act redundantly to promote aggregation behavior (de Bono et al., 2002). We conclude that QUI-1 acts in ASH, ADL, and potentially other neurons to downregulate O_2_-escape behavior.

**Figure 2.**
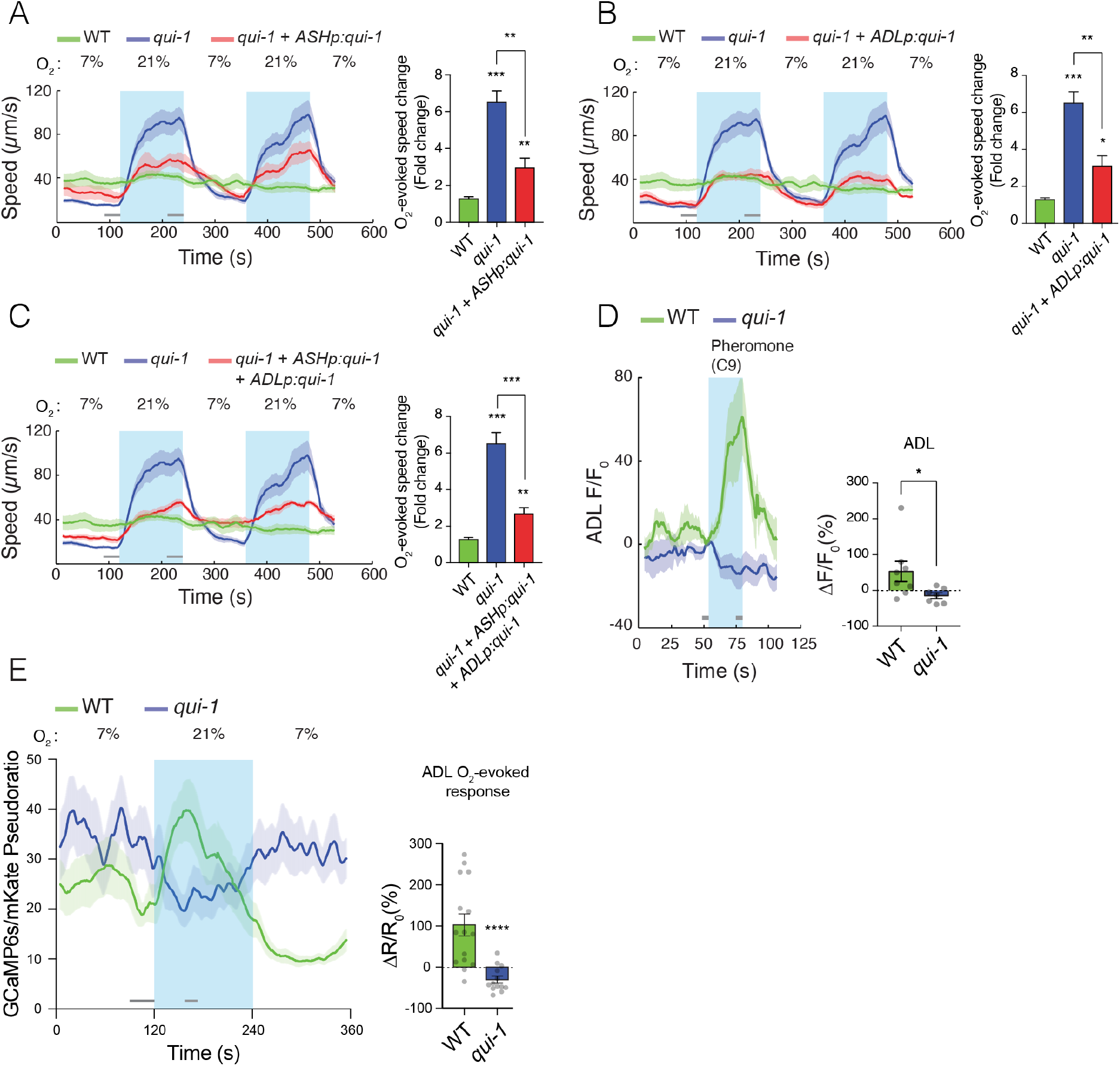
*qui-1* acts in ADL and ASH chemosensory neurons to inhibit O_2_-escape behavior, and it is required for pheromone-evoked Ca^2+^ responses in ADL. (**A-C**) Selective expression of *qui-1* in ASH (*sra-6p*), ADL (Δ*sre-1p*), or ASH + ADL (*sra-6p* + Δ*sre-1p*) neurons partially rescues the O_2_-escape phenotype. Left: Line shows average speed, while shading represents SEM. Grey bars show 30s time intervals used to average the animal’s speed. Right: Bar graph shows fold change in average speed at 21% O_2_ compared to 7% O_2_. N= 6–9 assays. (**D**) *qui-1* mutants lack pheromone-evoked Ca^2+^ responses in ADL. Left: Average GCaMP6s signal intensity (F) divided by baseline intensity (F_0_) is plotted over time. Shading shows SEM. Light blue rectangle indicates period of C9 pheromone stimulation. Right: Bar graph quantifying pheromone-evoked Ca^2+^ responses. ΔF/F_0_(%) was computed from 5s intervals before C9 stimulus was removed (F) and 5s before C9 stimulus was presented (F_0_), indicated by grey bars. N=8 (WT) and N=7 (*qui-1*). (**E**) *qui-1* mutants appear to lose O_2_-evoked Ca^2+^ responses in ADL. Left: Ca^2+^ levels reported as a pseudoratio between the GCaMP6s and mKate2 fluorescence signal. Both proteins are expressed under the ADL-specific promoter *srh-220p*. Grey horizontal bars show intervals (30s and 15s) used for calculating ΔR/R_0_ (%) in the Bar graph (right), which quantifies O_2_-evoked Ca^2+^ responses in ADL. N=15 (WT), N=13 (*qui-1*). Statistics: *, p value ≤0.05, **, p value ≤0.01, ***, p value ≤0.001, Mann-Whitney U test. Comparisons are with WT.

### QUI-1 is required for pheromone-evoked Ca^2+^ responses in ADL

*qui-1* mutants exhibit chemosensory response defects (Hilliard et al., 2004; Neal et al., 2016), but QUI-1’s role in these responses is not understood. Since O_2_-singlling reprograms the hub-and-spoke circuit, including ADL neurons (Fenk and de Bono, 2017), we speculated that disrupting *qui-1* alters ADL properties in a way that enhances circuit output in response to O_2_ stimuli. To probe how loss of *qui-1* alters ADL function, we first examined ADL responses to pheromones. In wild type control animals ADL neurons responded to the C9 ascaroside pheromone with a Ca^2+^ response, as expected (Jang et al., 2012), however this response was completely abolished in *qui-1* mutants (Figure 2D). This suggests that QUI-1 is required for sensory transduction of pheromone stimuli.
ADL neurons promote escape from 21% O_2_ (de Bono et al., 2002; Laurent et al., 2015). Consistent with this, ADL neurons exhibit a rise in Ca^2+^ in response to a 21% O_2_ stimulus. This ADL Ca^2+^ response depends on the URX neurons and the GCY-35/GCY-36 soluble guanylyl cyclases that are primary O_2_ sensors in these neurons (Fenk and de Bono, 2017; Zimmer et al., 2009). To investigate if disrupting *qui-1* alters O_2_-evoked Ca^2+^ responses in ADL, we imaged these responses using GCaMP6s, which provides improved sensitivity compared to indicators we used previously (Fenk and de Bono, 2017). Wild type animals exhibited a small but robust rise in Ca^2+^ levels upon stimulation with 21% O_2_, which rapidly returned to baseline (Figure 2E). This was followed by hyperpolarisation after the 21% O_2_ stimulus was removed (Figure 2E). Surprisingly, loss of *qui-1* abolished ADL O_2_-evoked Ca^2+^-responses, and stimulation with 21% O_2_ resulted, if anything, in ADL hyperpolarisation (Figure 2E). These data suggest that a simple increase in O_2_-evoked Ca^2+^ responses in ADL does not explain the increased ability of *qui-1* mutants to escape 21% O_2_.

### Disrupting *qui-1* enhances neurosecretion in ADL sensory neurons

Mutants lacking *qui-1* display defects in sensory perception while showing an augmented response to O_2_. This suggests altered cross-modal responses in the hub-and-spoke circuit might be responsible for this increase. Because ADL activity sustains O_2_-escape behavior, the *qui-1* behavioral phenotype could be explained by enhanced neurosecretion. To probe this, we used a fluorescently-tagged insulin-like peptide, DAF-28::mCherry, expressed specifically in ADL by the *srh-220* promoter. In *C. elegans*, insulin-like peptides are secreted through dense core vesicles and they accumulate in scavenger cells called coelomocytes; accumulation of fluorescently-tagged insulin-like peptides in these cells provides a readout of neurosecretion (Lee and Ashrafi, 2008). Using this assay, we found a striking increase in ADL neurosecretion in *qui-1* mutants compared to wild type (Figure 3A). Expressing wild type *qui-1* exclusively in ADL fully rescued this enhanced neurosecretion phenotype (Figure 3A). Increased insulin secretion levels cannot be explained by increased expression from the *srh-220* promoter: expression of free GFP from this promoter was not altered by mutations *qui-1* (Figure S3A). These data suggest that disrupting QUI-1 function in ADL enhances neurosecretion from this neuron.

**Figure 3.**
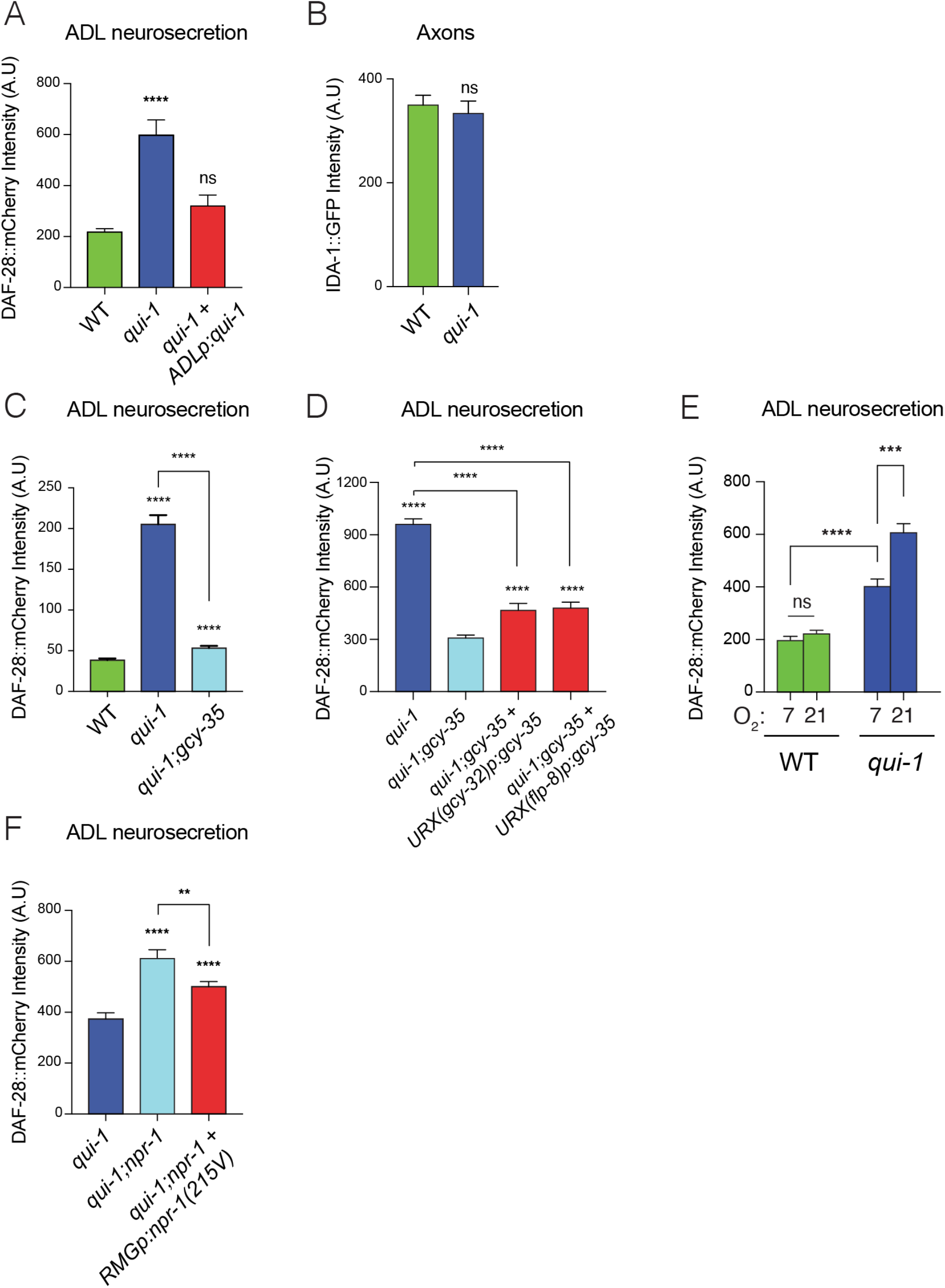
Loss of *qui-1* confers O_2_-evoked neurosecretion on ADL sensory neurons. (**A**) Disrupting *qui-1* increases neurosecretion from ADL. This phenotype is rescued by expressing *qui-1* cDNA specifically in ADL (Δ*sre-1p*). N=22 (WT), N=16 (*qui-1*), N=15 (*ADLp* rescue). (**B**) Loss of *qui-1* does not lead to increased axonal accumulation of DCVs. Bar graph show axonal levels of IDA-1::GFP, a DCV marker we expressed exclusively in ADL (*srh-220p*). N=42 (WT), N=34 (*qui-1*). (**C-D**) Increased ADL neurosecretion in *qui-1* mutants depends on the molecular O_2_-sensor *gcy-35* (**C**), which acts partly in URX neurons (**D**). (C) N=21 (WT), N=30 (*qui-1*), N=27 *(qui-1;gcy-35*). (D) N=60 (*qui-1*), N=46 (*qui-1;gcy-35*), N=46 (URX *gcy-32p* rescue), N=32 (URX *flp-8p* rescue). (**E**) O_2_ levels modulate ADL neurosecretion in *qui-1* mutants but not WT. Animals were raised at either 7% or 21% O_2_ from egg to young adult. N=30 (WT 7%), N=22 (WT 21%), N=23 (*qui-1* 7%), N=35 (*qui-1* 21%). (**F**) RMG signalling contributes to ADL neurosecretion in *qui-1* mutants. Rescue was achieved using two promoters overlapping only in RMG. *flp-21* promoter drives a floxed transcriptional STOP signal followed by *npr-1*(215V) isoform (*flp-21p:flox:STOP:flox:npr-1(215V)*) while *ncs-1* promoter drives the Cre recombinase (*ncs-1p:Cre*). N=27 (*qui-1*), N=22 (*qui-1;npr-1*), N=33 (*RMGp* rescue). In **A, C, D, E, F** bar graphs report the accumulation of DAF-28::mCherry fluorescence in coelomocytes following its release from ADL. Statistics: **, p value ≤0.01, ***, p value ≤0.001, ****,p value ≤0.0001, ns=not significant. Mann-Whitney U test. Comparisons are against WT in A, B and C, against *qui-1;gcy-35* in D and against *qui-1* in F.

Increased neurosecretion could reflect delivery of a larger number of dense core vesicles (DCVs) to release sites. To ask if *qui-1* alters DCV trafficking, we tagged IDA-1, a DCV-associated protein, with GFP and expressed this fusion protein exclusively in ADL. ADL is highly polarized: its cell body projects a dendrite anteriorly, to the animal’s nose and an axon that splits at the nerve ring into ventral and dorsal projections that form synapses with post-synaptic partners. As expected, IDA-1::GFP fluorescence was localized to small bright puncta along ADL axons and more diffusely in the ADL cell body (Figure S3C II-IV). No signal was detected in dendrites. To quantify possible differences, we measured the intensity of IDA-1::GFP signal. *qui-1* mutants did not show gross differences in the axonal distribution of IDA-1::GFP (Figure 3B). To assess if more IDA-1::GFP was retained in the cell body in *qui-1* mutants, we compared fluorescence signals between *qui-*1 and wild type but did not observe any differences (Figure S3B). Moreover, *qui-1* mutants did not show obviously altered ADL morphology (Figure S3C I-III). These data suggest that enhanced neurosecretion from ADL in *qui-1* mutants is not due to increased DCVs accumulation in axons but may reflect an increased rate of release.

### Absence of *qui-1* increases neurosecretion from ADL in response to O_2_-circuit input

Why do *qui-1* mutants exhibit increased neurosecretion from ADL neurons? A simple hypothesis, prompted by the increased behavioral response of *qui-1* mutants to 21% O_2_, is that enhanced ADL neurosecretion is due to stronger coupling to input from URX. The soluble guanylyl cyclase GCY-35 acts as the main oxygen molecular sensor: null mutations in *gcy-35* disrupt O_2_-evoked responses both at the circuit and behavioral level (Busch et al., 2012; Laurent et al., 2015; Zimmer et al., 2009). Consistent with this, disrupting *gcy-35* almost completely abolished the enhanced neurosecretion of *qui-1* mutants (Figure 3C). Over-expressing wild type GCY-35 in URX, using the *flp-8* or *gcy-32* promoters, rescued the ADL neurosecretion phenotype of *qui-1*;*gcy-35* double mutants, although not completely (Figure 3D). We conclude that increased neurosecretion from ADL neurons in *qui-1* mutants reflects an enhanced response to O_2_ partly mediated by URX neurons.

Our experiments with *qui-1*;*gcy-35* double mutants predict that manipulating ambient O_2_ levels should shape ADL neurosecretion in *qui-1* mutants. To investigate this hypothesis, we grew wild type and *qui-1* mutants at 7% and 21% O_2_ and assayed neurosecretion from ADL. In wild type animals ADL neurosecretion was unaffected by O_2_ experience (Figure 3E). By contrast, neurosecretion from ADL was significantly modulated by O_2_ experience in *qui-1* mutants (Figure 3E). Mutants kept at low O_2_ concentrations showed markedly less ADL neurosecretion than animals kept at 21% O_2_ (Figure 3E). Together, these data support the hypothesis that disrupting *qui-1* confers O_2_-evoked neurosecretion on ADL neurons.

URX and ADL neurons are connected by gap junctions to RMG interneurons in the hub-and-spoke circuit (Cook et al., 2019; Macosko et al., 2009). Signalling from the NPR-1 neuropeptide receptor in RMG modulates communication across the hub-and-spoke circuit. In the N2 genetic background, a hyperactive version of this neuropeptide receptor, NPR-1 (215V), impedes communication across the circuit (Macosko et al., 2009). To test if RMG activity alters ADL neurosecretion, we assayed *qui-1* and *qui-1;npr-1* double mutants. We observed higher levels of neurosecretion from ADL in *qui-1;npr-1* double mutants (Figure 3F). Over-expressing NPR-1 215V in RMG partially rescued the ADL phenotype of *qui-1*;*npr-1* double mutants (Figure 3F). We conclude that NPR-1 signalling in RMG neurons can suppress neurosecretion from ADL. Taken together, these and previous data suggest enhanced ADL neurosecretion in *qui-1* mutants is principally driven by increased ADL responsiveness to O_2_ input from the hub-and-spoke circuit.

### Disrupting sensory perception confers O_2_-evoked neurosecretion on ADL sensory neurons

Is the increased coupling of ADL to the hub-and-spoke circuit specific to *qui-1* mutants or an adaptation to impaired sensory perception? A group of genes involved in sensory perception and associated with Bardet-Biedl syndrome, called *bbs* genes in *C. elegans*, has been proposed to reduce, by an unknown mechanism, neurosecretion (Lee et al., 2011). *bbs* genes encode components of a large protein complex involved in intraflagellar transport, the BBsome, which binds cargo vesicles to motor proteins for delivery to cilia. *bbs* mutants exhibit sensory defects, and, like *qui-1*, show increased dense-core vesicle release from ADL (Lee et al., 2011). We asked if *bbs* mutants also show increased O_2_-escape behavior. Of the five *bbs* mutants we studied, three, *bbs-7, bbs-1*, and *bbs-2*, showed *qui-1*-like O_2_-escape behavior (Figure 4A and Figure S4A to C). These data suggest more sensory defective mutants could display elevated O_2_-escape behavior.

**Figure 4.**
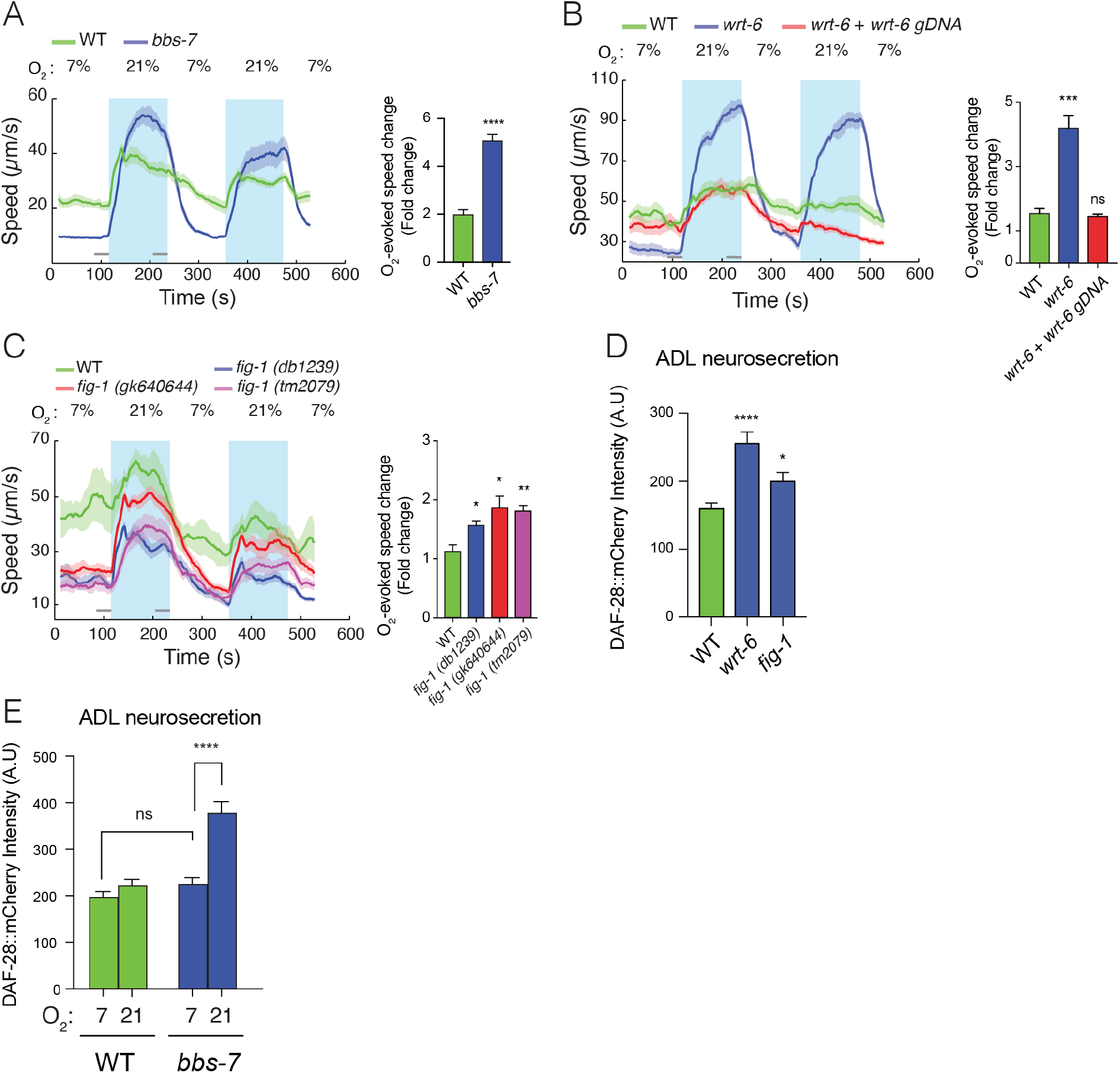
A range of mutants with sensory defects exhibit increased behavioral responsiveness to O_2_ and confer O_2_-evoked neurosecretion on ADL sensory neurons. (**A**) *bbs-7* mutants, which lack a subunit of the BBSome complex and have impaired cilia formation and function, show increased O_2_-escape behavior (See also Figure S4A-C). (**B**) *wrt-6* mutants, which are defective in a hedgehog-related gene expressed in glia surrounding the sensory endings of chemosensory neurons, show increased O_2_-escape behavior. The phenotype is rescued by a wild type copy of *wrt-6* (**C**) Null mutants of *fig-1*, another gene expressed in glia whose loss causes chemosensory defects, show increased O_2_-escape behavior similar to the *fig-1(db1239)* allele isolated in our screen (See Figure S4E). (**A-C**) Left: Lines show average speed, shading represents SEM and grey bars represent 30s intervals used to average the animal’s speed. Right: Bar graphs show fold change in average speed at 21% O_2_ compared to 7% O_2_. N=6-9 assays. (**D**) *wrt-6* and *fig-1* mutants show increased neurosecretion from ADL. N=22 (WT), N=21 (*wrt-6*), N=18 (*fig-1*). (**E**) Increased ADL neurosecretion in *bbs-7* mutants reflects increased responsiveness to O_2_ stimuli. Animals experienced 7% or 21% O_2_ from egg to young adult, as indicated. N=30 (WT 7%), N=22 (WT 21%), N=25 (*bbs-7* 7%), N=25 (*bbs-7* 21%). (**D-E**) Bar graphs show DAF-28::mCherry fluorescence taken up by coelomocytes following release from ADL. Statistics: *, p value ≤0.05, **, p value ≤0.01, ***, p value ≤0.001, ****, p value ≤0.0001, ns=not significant, Mann-Whitney U test. Unless indicated, comparisons are against WT.

A search of the sequencing data from our mutant collection revealed two mutant strains carried missense mutations in genes previously associated with impaired sensory perception, *wrt-6* (WaRThog, a hedgehog-related protein) and *fig-1* (dye-*F*illing abnormal, expressed *I*n *G*lia) (Bacaj et al., 2008; Hao et al., 2006). The *wrt-6* (*db102*) allele substituted a conserved threonine residue (T460I) (Figure S4D) essential for autocleavage and activation of Hedgehog-like secreted proteins; the *fig-1* (*db1239*) allele changed a cysteine in a C6 domain into a tyrosine (C1951Y) (Figure S4E). A wild type copy of *wrt-6* entirely rescued the O_2_- escape phenotype of *db102* mutants, confirming that these phenotypes reflect loss of *wrt-6* function (Figure 4B). To test if defects in *fig-1* elevated O_2_-escape behavior, we assayed multiple *fig-1* loss-of-function alleles (Figure S4E). All *fig-1* alleles showed increased O_2_- responses (Figure 4C), confirming that the absence of *fig-1* leads to stronger O_2_-escape behavior. We also injected *fig-1* mutants with a wild type copy of *fig-1* but failed to rescue O_2_- escape behaviour (data not shown). Appropriate protein levels may be necessary for correct FIG-1 function.

Together, our data suggest that compromising sensory input increases ADL’s responsiveness to O_2_ input from the hub-and-spoke circuit. To test this, we measured ADL neurosecretion in *wrt-6* and *fig-1* mutants and observed a robust increase in both mutants (Figure 4D). *wrt-6* and *fig-1* are expressed in glia and not neurons (Bacaj et al., 2008; Hao et al., 2006) and are unlikely to regulate neurosecretion directly. We next asked if enhanced neurosecretion from ADL in sensory defective mutants depended on O_2_ input. We raised *bbs-7* mutants, which showed the strongest O_2_-escape behavior among the sensory defective mutants we had studied, at 7% and 21% O_2_ and measured ADL neurosecretion. *bbs-7* mutants grown at 7% O_2_, when URX–RMG activity is low, lost their enhanced neurosecretion phenotype and showed secretion levels indistinguishable from wild type animals reared at 7% O_2_ (Figure 4E). These data suggest that ADL neurons release more DCVs in *bbs, wrt-6*, and *fig-1* mutants than wild type animals in response to input from the O_2_ circuit.

### Elevating NPR-22 expression in ADL underpin O_2_-evoked neurosecretion

Defects in sensory perception reprogram ADL properties to enhance neurosecretion in response to input from URX–RMG. To investigate the molecular details behind this reprogramming, we labelled ADL neurons by expressing mKate2 using an ADL-specific promoter (*srh-220*p), used fluorescence-activated cell sorting (FACS) to sort ADL from freshly dissociated wild type and *qui-1* mutants and then profiled the ADL transcriptome using RNA sequencing (RNAseq). Enrichment analysis highlighted ADL as the most enriched neural class in our data set (Figure S5A). Our RNAseq data also included known ADL-specific transcripts such as *srh-234* (Gruner et al., 2014) and *srh-279* (Vidal et al., 2018), and *qui-1* itself, but not transcripts expressed in neighbouring neurons such as ASK and ASI (Supplementary File 1 and data not shown). Consistent with its function as a chemosensory neuron, ADL expresses a large number of chemoreceptors (Supplementary File 2), several neuropeptide receptors (Supplementary File 3) and high levels of neuropeptides (Supplementary File 4).
To identify gene involved in ADL’s reprogramming, we next examined genes differentially regulated between wild types and *qui-1* mutants in ADL (Supplementary File 5). Principal component analysis (PCA) confirmed we could robustly differentiate *qui-1* from control samples (Figure S5B) while also highlighting how much ADL’s transcriptional landscape is altered by loss of *qui-1*; the majority of differentially regulated genes are strongly upregulated in mutant samples (Figure S5C and S5D). *qui-1* mutants display a large increase in ADL neurosecretion. Consistent with this, when we selected all known genes associated with DCV release (Hobert, 2006) that were also differentially regulated, almost all them were elevated in *qui-1* mutants (Figure 5A). This is likely to sustain the higher rate of neurosecretion. The absence of *qui-1* also reprogramed chemosensory receptor levels: more than half of all the chemoreceptors expressed in ADL were differentially regulated (Figure S5E and S5F). We conclude that loss of *qui-1* largely reprograms ADL properties and enhances genes associated with synaptic release.

**Figure 5.**
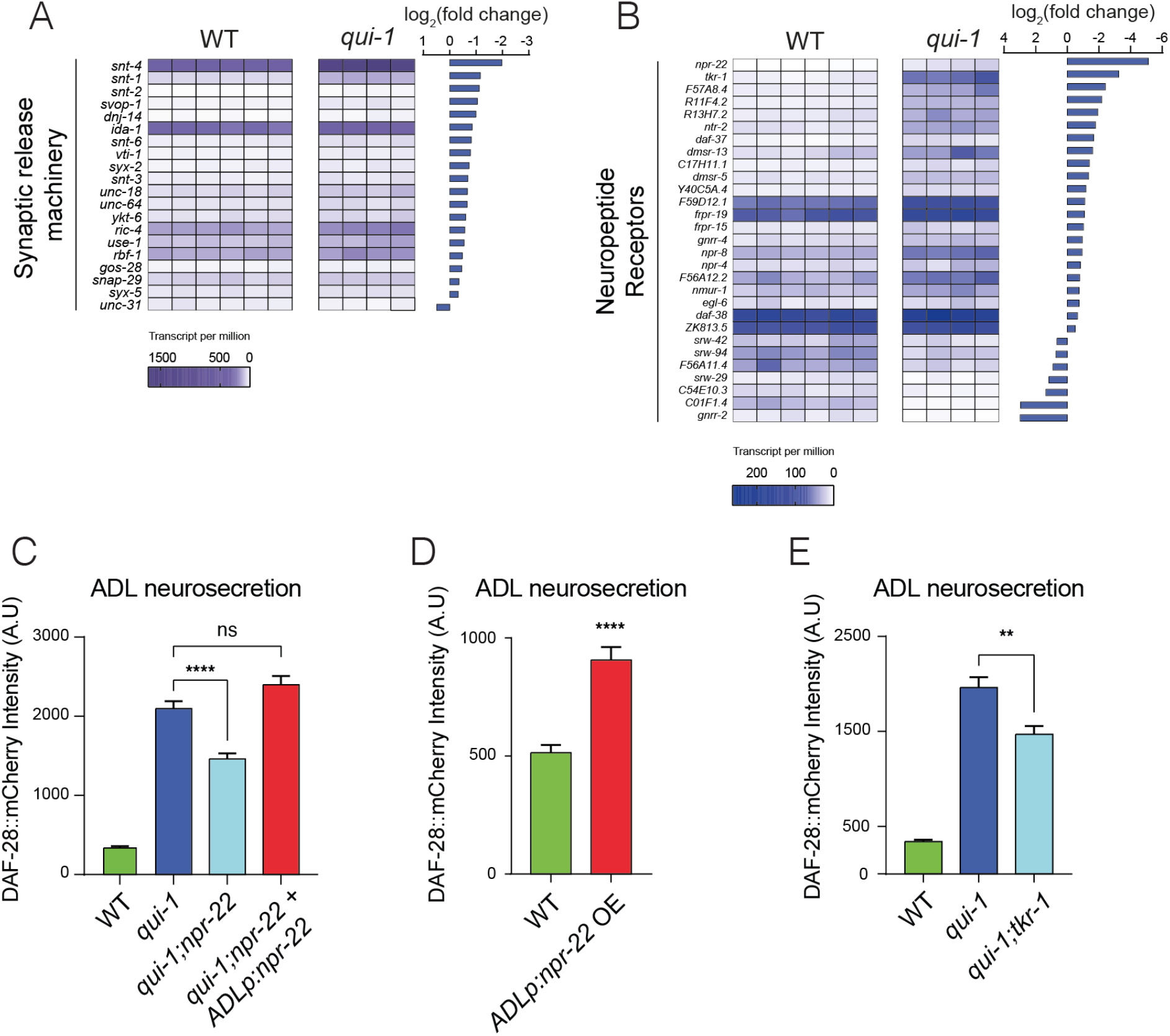
Increased O_2_-evoked neurosecretion from ADL in *qui-1* mutants is associated with reprogrammed peptidergic communication. (**A**) *qui-1* mutants upregulate a suite of genes that control synaptic and/or DCV release. (**B)** Loss of *qui-1* reprograms neuropeptide receptor expression in ADL. (**A-B**) Heat maps show expression values (transcript per million, tpm) for each gene across biological replicates, indicated by rows. To target analysis, we used lists generated in a review of the neural genome (Hobert, 2006), and selected the annotation “Synaptic release machinery” (**A**) or “Neuropeptide receptor” (**B**). All genes matching these criteria were included unless their expression was below 10 tpm in both genotypes or the q-value was >0.05 (See Supplementary File 5). (**C**) The neuropeptide receptor NPR-22 promotes neurosecretion from ADL in *qui-1* mutants; this phenotype can be rescued by expressing *npr-22* cDNA (encoding NPR-22 isoform b) from an ADL-specific promoter (*srh-220*p). N=33 (WT), N=54 (*qui-1*), N=53 (*qui-1;npr-22*), N=52 (*ADLp* rescue). (**D**) Overexpressing the same *npr-22* cDNA construct specifically in ADL is sufficient to stimulate neurosecretion in WT. N=46 (WT), N=44 (*ADLp* OE). (**E**) The tachykinin receptor *tkr-1* also stimulates ADL neurosecretion in *qui-1* mutants. N=38 (WT), N=37 (*qui-1*), N=38 (*qui-1;tkr-1*). (**C–E**) Bar graphs show intensity of DAF-28::mCherry accumulated in coelomocytes following released from ADL. Statistics: **, p value ≤0.01, ****, p value ≤0.0001, ns, not significant, Mann-Whitney U test. In C and E comparisons are against *qui-1*, while in D comparisons are against WT.

Our data indicated that in *qui-1* and other sensory-defective mutants ADL responds more strongly to input from O_2_-sensing neurons by releasing more DCVs. Cyclic adenosine monophosphate (cAMP), a common second messenger for the activation of neuropeptide receptors, strongly stimulate DCVs release (Costa et al., 2017). This together with absence of increased O_2_-evoked Ca^2+^ responses in ADL in *qui-1* mutants (Figure 2E) disfavored increased synaptic communication between ADL and the hub-and-spoke circuit. Instead, we hypothesized that ADL reprogramming in sensory-defective mutants involves an altered response to neuropeptide input. Several neuropeptide receptors were differentially expressed in ADL between wild type and *qui-1* (Figure 5B). Most prominent of these was neuropeptide receptor 22 (*npr-22*), one of the most highly upregulated genes in *qui-1* overall (Figure S5D). To investigate if elevated NPR-22 levels in *qui-1* mutants explained the increased DCV release in ADL, we compared ADL neurosecretion in *qui-1* and *qui-1;npr-22* animals. The double mutants showed robustly reduced neurosecretion compared to *qui-1* mutants (Figure 5C). This phenotype was completely rescued by expressing *npr-22* in ADL, confirming that *npr-22* is necessary to sustain ADL’s higher neurosecretion levels. *npr-22* does not seem to be expressed at appreciable levels in wild type ADL according to our profiling data (Supplementary File 5). To test if inducing *npr-22* expression is sufficient to stimulate ADL neurosecretion, we over-expressed the neuropeptide receptor in wild type ADL neurons. Increasing *npr-22* expression was sufficient to induce a higher rate of neurosecretion from ADL (Figure 5D). Thus, increasing peptidergic signalling through NPR-22 receptors is necessary and sufficient to promote ADL O_2_-evoked neurosecretion.

Disrupting *npr-22* did not completely suppress the neurosecretion phenotype of *qui-1* mutants. We investigated if other neuropeptide receptors whose expression in ADL is induced in *qui-1* mutants augmented O_2_-evoked neurosecretion from ADL. The second neuropeptide receptor gene in our list (Figure 5B) was the TachyKinin Receptor 1, *tkr-1*. Loss of *tkr-1* robustly decreased neurosecretion from ADL to levels comparable to those in *qui-1;npr-22* (Figure 5E). We conclude that disrupting sensory responsiveness in ADL, by the *qui-1* mutation, increases the coupling of this neuron to the hub-and-spoke circuit by inducing expression of neuropeptide receptors that enhance DCV release from ADL in response to pre-synaptic input.

## Discussion

Cross-modal plasticity is thought to involve recruitment of impaired neurons to process additional sensory modalities. The molecular details of such rearrangement are not yet clear. Here, we use forward genetics as our entry point to sought mechanisms that increase *C. elegans’* responsiveness to an oxygen (O_2_) sensory cue. Multiple mutants we identify simultaneously increase responsiveness to O_2_ while disrupting other sensory responses – a hallmark of cross-modal plasticity. We analyse one of these mutants, *qui-1*, which is defective in the ortholog of mammalian NWD1, in depth. We show that loss of QUI-1 prevents the ADL sensory neurons from responding to pheromone, but increases ADL neurosecretion in response to input from upstream O_2_-sensing neurons. Loss of *qui-1* thus recruits ADL sensory neurons more strongly into the O_2_-sensing circuit. We observe a similar role change in animals defective in Bardet-Biedel syndrome (*bbs*) proteins that mediate intraciliary transport in sensory neurons. Loss of *qui-1* reprograms gene expression in ADL neurons and induces expression of neuropeptide receptors, including NPR-22 and TKR-1. Upregulation of these neuropeptide receptors support the coupling of ADL neurosecretion to O_2_ input in *qui-1* mutants. We propose that impairing sensory perception can sensitise sensory neurons to other sensory modalities by reprogramming peptidergic circuits.

ADL-specific expression of *qui-1* rescues both the increased O_2_-escape phenotype of *qui-1* mutants and enhanced neurosecretion from ADL (Figure 2B and 3A), indicating that *qui-1* acts cell-autonomously to regulate neurosecretion. Impairing the primary O_2_-sensing mechanism, by disrupting the molecular oxygen sensor GCY-35, restores ADL neurosecretion in *qui-1* mutants to levels observed in wild controls (Figure 3C), confirming that increased ADL neurosecretion in *qui-1* is driven primarily by activity originating outside ADL. Together with data showing that *qui-1* mutants exhibit normal O_2_-evoked Ca^2+^ responses in URX and RMG neurons (Figure S1C and S1D), this suggests that in *qui-1* mutants ADL neurons are more sensitive to incoming pre-synaptic activity.

Selectively expressing *gcy-35* cDNA in the URX O_2_-sensor partially rescues the ADL neurosecretion phenotype of *qui-1*;*gcy-35* mutants (Figure 3D), confirming that URX helps drive increased ADL neurosecretion in *qui-1* mutants. Disrupting the inhibitory neuropeptide receptor *npr-1* further increases ADL neurosecretion in *qui-1* mutants, and this phenotype is partially rescued by expressing *npr-1* cDNA specifically in RMG interneurons (Figure 3F). This adds further support to our model, and indicates that activity from the hub-and-spoke circuit propagates from URX–RMG to ADL to stimulate neurosecretion. Consistent with this, in *qui-1* mutants, but not in wild type, ADL neurons show O_2_-evoked neurosecretion: prolonged exposure to low (7%) or high (21%) O_2_ concentrations modulates ADL neurosecretion (Figure 3E). These data confirm how disrupting *qui-1* recruits ADL more strongly into the rest of the hub-and-spoke circuit. They also suggest a model to explain why *qui-1* mutants show increased O_2_-escape behaviour: the absence of *qui-1* sensitises ADL to incoming activity from O_2_-sensors and increases ADL neurosecretion which is likely to stimulate O_2_-escape behaviour.

Cross-modal plasticity has been described previously in *C. elegans*, in a touch receptor/olfactory circuit paradigm (Rabinowitch et al., 2016), which is mediated by an alternative mechanism from the one we describe here. Worms with touch receptor defects show enhanced odorant responses compared to wild type because activated touch receptors release an inhibitory neuropeptide, FLP-20, that downregulates communication between the AWC olfactory neurons and their post-synaptic target, the AIY interneurons. Loss of touch receptor function thus enhances odorant responses by disinhibiting AWC–AIY communication. In this mechanism the defective touch receptors do not to contribute to the enhanced odorant sensing, a contrast to the mechanism we describe. FLP-20 appears to act as a general arousal signal, relaying information about mechanical stimulation to multiple circuits (Chew et al., 2018).

Several questions remain outstanding. How does disrupting *qui-1*, and ADL sensory function, lead to reprogramming of ADL gene expression? Comparing gene expression in ADL between *qui-1* and wild type controls reveals altered expression of several transcription factors, including members of the nuclear hormone receptor family (*nhr*), the *egl-46* zinc-finger protein (Wu et al., 2001) and the storkhead box protein *ham-1* (Feng et al., 2013). Some of these transcription factors show substantial (e.g. >30 fold) induction, and are orthologs of immediate early genes in mammals. These transcription factors may contribute to reprogramming ADL.

Previous work has shown that mutations in BBS-7, a conserved protein involved in trafficking of molecular cargos along the primary cilium of neurons, and linked to Bardet-Biedl syndrome (Liu and Lechtreck, 2018; Tan et al., 2007), lead to increased neurosecretion from ADL (Lee et al., 2011). Mutations in *bbs-7* cause defects in cilia formation and in sensory perception. Our data suggest that increased neurosecretion from ADL in *bbs* mutants may reflect increased coupling to the O_2_ circuit (Figure 4E). Interestingly, *bbs-7* mutants show a range of other physiological phenotypes including small body size and delays in development (Lee et al., 2011; Mok et al., 2011). Both of these phenotypes can be suppressed by mutating *gcy-35* (Mok et al., 2011), consistent with them resulting from enhanced responses to O_2_. Other genes associated with intraflagellar transport (IFT) in sensory cilia function have been shown to promote aggregation, both in *C. elegans* (de Bono et al., 2002; Moreno and Sommer, 2018) and *P. pacificus* (Moreno et al., 2017), a nematode species distantly related to *C. elegans*. We speculate that these IFT mutants, like the *bbs* mutants and *qui-1*, enhance O_2_ escape behavior by increasing the coupling of sensory-defective sensory neurons to input from the O_2_ circuit.

Conceptually, our findings resonate with studies in vertebrates which find that loss of a sensory modality can lead to recruitment of input-deprived sensory cortex to process information from spared senses (Lee and Whitt, 2015; Petrus et al., 2014; Rauschecker, 1995). In the nervous system of *C. elegans*, the defective ADL sensory neurons becomes sensitized to pre-synaptic input associated with a different modality, O_2_ sensing. This recruitment is supported by a transcriptional program in the sensory defective ADL that induces the expression of neuropeptide receptors including NPR-22 and TKR-1 (Figure 6A). We believe our data support a conserved role for reconfiguring peptidergic circuits in support of cross-modal recruitment.

**Figure 6.**
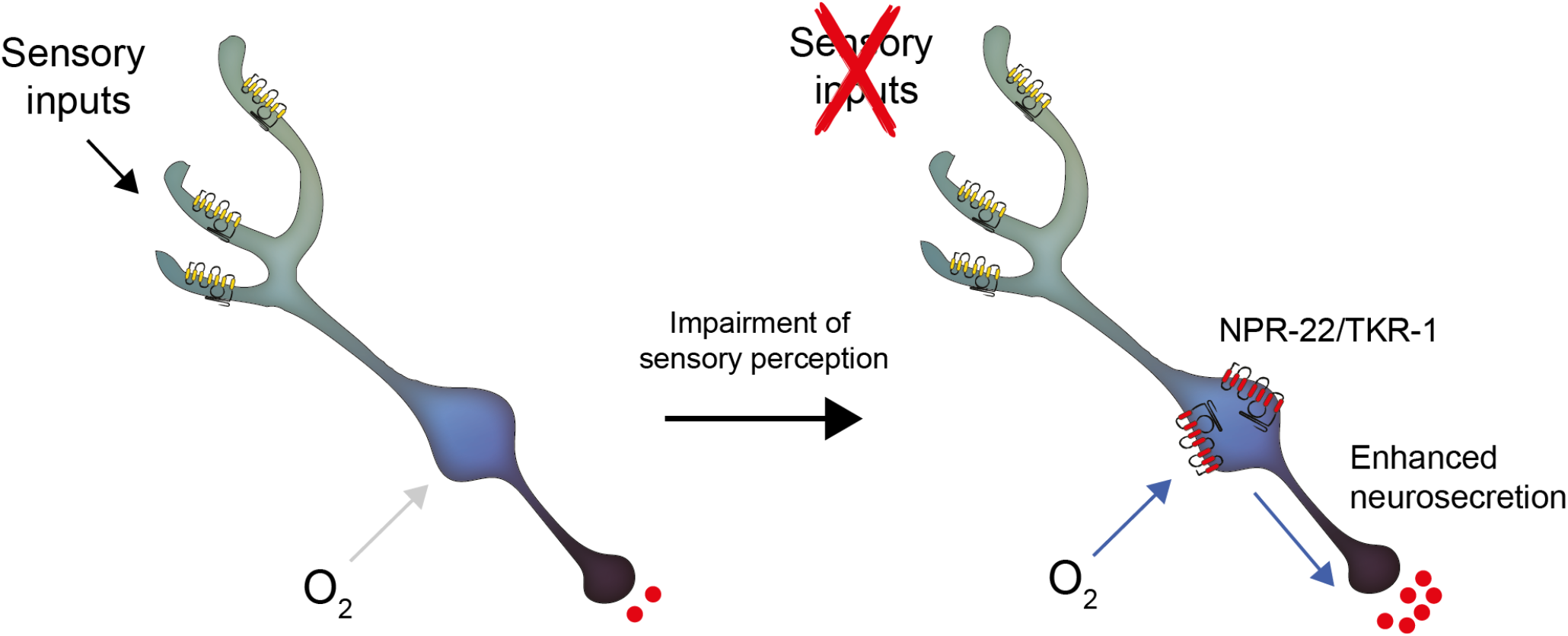
Model depicting the effect of disrupting sensory perception on ADL sensory neurons. Schematic depiction of model explaining ADL O_2_-evoked neurosecretion in sensory defective mutants. When ADL sensory perception is functioning (left side), O_2_ does not modulates its neurosecretion, despite ADL receiving signals from O_2_-sensors. When sensory perception is impaired (right side) ADL undertakes a transcriptional reprogramming resulting in stronger coupling of ADL neurosecretion to input from O_2_ sensors. Much of the increase in O_2_-evoked neurosecretion is conferred by increased expression of two receptors, NPR-22 and TKR-1, in ADL itself.

## Material and Methods

### Strains

*C. elegans* were grown at room temperature under standard conditions (Brenner, 1974). All assays used young adults (< 24 hours old). Transgenic animals were obtained by injecting DNA mixtures of an expression construct or fosmid and a co-injection marker, each at 20 – 40ng/μL. A list of strains used is provided in Supplementary Information. *E. coli* OP50 cultures were grown in 2xTY broth and used to seed NGM plates.

### Mutagenesis

Animals were mutagenized with a 50 mM solution of ethylmethane sulfonate (EMS) in M9 buffer (Brenner, 1974). To isolate mutants that preferentially aggregated on thick food we placed an ∼ 0.2 x 0.2 cm patch of concentrated *E. coli* OP50 at the centre of a much thinner circular lawn of OP50, ∼5 cm in diameter, seeded on a 9 cm NGM dish. F2 progeny of mutagenized animals were washed 2x in M9 buffer and kept without food for ∼30 minutes before being pipetted outside the thin bacterial lawn. Test experiments showed that under these conditions animals from non-aggregating strains strongly inhibited movement upon encountering the thin lawn. By contrast, individuals from aggregating strains continued moving quickly on the thin lawn but settled when they encountered the thick bacterial patch (Figure 1B – Selection 1). Potential aggregating mutants were collected from the thick bacterial patch ∼ 60 minutes after animals were added on the plate. These animals were then individually placed on a seeded NGM plate and their progeny scored for aggregation behavior.

### Behavioral assays

#### Aggregation assays

The assay was performed as described (de Bono and Bargmann, 1998). 60 young adults were picked onto assay plates seeded 2 days earlier with 200 μL OP50. Animals were left undisturbed for 3 hours and scored as aggregating if they were in a group of 3 or more individuals in contact over > 50% of their body length. Animals were considered to be at the lawn border if they were within 2 mm of the lawn edge. Scoring was performed blind to genotype. For each biological replicate:

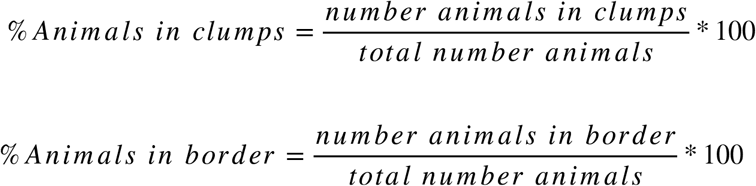

#### Locomotion assays

Assays were performed as described previously (Laurent et al., 2015). Low peptone NGM plates (0.13% w/v bactopeptone) were seeded with 60 μL of OP50 broth 2 days before the assay. On the day of the assay, test plates were prepared by cutting the edge of the bacterial lawn using a rubber stamp. Around 20 young adults were picked onto the lawn and left undisturbed for 10 minutes before starting the assay. A PDMS chamber was placed on top of the bacterial lawn and defined gas mixtures delivered to the chamber at 1.25 mL/min using a pump (PHD 2000, Harvard Apparatus). Worms were allowed to adapt to 7% O_2_ for 2 minutes before videorecording started. Worms were recorded for 9 minutes while the gas mixture pumped into the chamber was changed from 7% O_2_ to 21% O_2_ every 2 minutes. Videorecordings were acquired at 2 frames per second (fps) using a Grasshopper camera (Point Grey) mounted on a stereomicroscope (Leica MZ6 and MZ7.5). Videos were analysed and animal speed calculated using a custom-written MATLAB software (Zentracker: https://github.com/wormtracker/zentracker). Average speed values were extracted using Metaverage, a custom-written MATLAB software. Bar graphs show a ratio of the average speed 30s before the end of the first 21% O_2_ stimulus and the average speed 30s before O_2_ levels were first switched from 7% O_2_ to 21% O_2_. This ratio was computed for all biological replicates (independent assays) and entered into Prism for statistical analysis.

### Molecular Biology

#### DNA extraction for whole-genome sequencing

Genomic DNA for whole-genome sequencing was isolated from 5-10 crowded 5 cm NGM plates. Animals were washed off plates in M9 buffer, rinsed 2x in M9 buffer to remove OP50, and frozen at -80°C. Genomic DNA was extracted from thawed samples using the DNeasy Blood and Tissue Kit (Qiagen). Samples were left in Lysis buffer (Buffer AL) for 3h at 56°C and DNA isolated following manufacturer’s instructions.

#### Library preparation

Libraries for whole-genome sequencing were prepared using Nextera XT DNA Library kit (Illumina) following the manufacturer’s instructions. Library quality was checked on a Bioanalyzer using Agilent High Sensitivity Gel. Library concentration was assessed using KAPA Library Quantification Kits for Illumina (KAPA Biosystems) prior to sequencing on the Illumina HiSeq 4000 platform.
RNAseq of identified neurons was adapted from Picelli *et al*.,. Briefly, fluorescently-labelled neurons collected by FACS were lysed in 10 μL of 0.2% Triton X-100 (vol/vol) and 2U/μL RNase inhibitors. The reverse transcription reaction volumes were adjusted to a 10 μL input and cDNA prepared using oligo dT primers and template-switching oligos (TSO) to enrich for polyadenylated transcripts and allow for pre-amplification of cDNA. cDNA was pre-amplified using custom PCR primers. 50 μL of PCR product from the pre-amplification step were purified using 50 μL of Ampure XP beads (Beckman Coulter), resuspended, and used at a concentration of 0.2 ng/uL as input for library preparation using the Nextera XT DNA Kit. Library preparation followed the manufacturer’s instruction. The quality of RNAseq libraries was assessed on a Bioanalyzer (Agilent) using High Sensitivity Gels (Agilent). Library concentration was measured using a Qubit dsDNA High Sensitivity Kit (Thermo Fisher Scientific) and sequenced on the Illumina HiSeq 4000 platform.

### Ca^2+^ imaging

#### Ca^2+^ imaging of O_2_-evoked URX activity

5 – 10 young adult animals (< 24 hrs old) expressing the YC2.60 Ca^2+^ sensor were glued to agarose pads (2% in M9 buffer, 1 mM CaCl2) using Dermabond tissue adhesive, with their body immersed in OP50 washed off from a seeded plate using M9. The animals were quickly covered with a PDMS microfluidic chamber and 7% O_2_ pumped into the chamber for 2 min before imaging, to allow animals to adjust to the new conditions. Neural activity was recorded for 6 minutes with switches in O_2_ concentration every 2 minutes. Imaging was on an AZ100 microscope (Nikon) equipped with a TwinCam adaptor (Cairn Research), two ORCA-Flash4.0 V2 digital cameras (Hamamatsu), and an AZ Plan Fluor 2× objective with 2× zoom. Recordings were at 2 fps with a 500 ms exposure time. Excitation light from a C-HGFI Intensilight lamp (Nikon) was passed through a 438/24 nm filter and an FF458-DiO2 dichroic (Semrock). Emitted light was passed to a DC/T510LPXRXT-Uf2 dichroic filter in the TwinCam adaptor cube and then through 483/32 nm (CFP) or 542/27 nm (YFP) filters before collection on the cameras.

#### Ca^2+^ imaging of O_2_-evoked RMG activity

The imaging protocol was performed as reported for URX, except that *db104* mutants were imaged on an Axiovert 200 microscope (Zeiss) with a 40x NA 1.2 C-Apochromat objective using an EMCCD Evolve 512 Delta camera (Photometrics), which gave higher signal-to-noise than the Nikon AZ100.

#### Ca^2+^ imaging of pheromone-evoked ADL activity

We used olfactory chips (Microkosmos LLC, Michigan, USA) to image young adults (< 24 hrs old) expressing the GCaMP3 Ca^2+^ sensor specifically in ADL, as previously described (Chronis et al., 2007; Jang et al., 2012). Animals were kept under a constant flow of M13 buffer and after 2 min stimulated for 20 sec with C9 pheromone (10 nM in M13 buffer). Ca^2+^ imaging used a 40x NA 1.2 C-Apochromat lens on an Axiovert 200 microscope (Zeiss) equipped with Dual View emission splitter (Photometrics) and a EMCCD Evolve 512 Delta camera (Photometrics). Acquisition was at 2 fps with a 100 ms exposure. Excitation light was from a Lambda DG-4 (Sutter Instruments) and it was passed through an excitation filter (AmCyan,Chroma), a dichroic filter for GCaMP and RFP. A beam splitter (Optical Insights) was used to separate the GCaMP and RFP signal using a dichroic filter 514/30-25 nm (GFP) and 641 nm (RFP) (Semrock).

#### Calcium imaging of O_2_-evoked ADL activity

We imaged young adults (< 24 hrs old) co-expressing GCaMP6s and mKate2 from the ADL-specific *srh-220p*, in a bi-cistronic construct. Animals were immobilised with Dermabond glue, placed under a PDMS chamber and imaged on the same microscope and imaging set-up used to image pheromone-evoked Ca^2+^ in ADL. Prior to recording activity, animals were pre-stimulated for 3 minutes to extinguish light-evoked ADL responses. Acquisition was at 2 fps with a 100 ms exposure. O_2_ concentration was switched between 7% and 21% O_2_ every 2 minutes.

All recordings were analysed using Neuron Analyser, a custom written MATLAB program available at https://github.com/neuronanalyser/neuronanalyser.

#### Analysis

Bar graphs showing 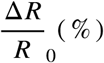 used YFP/CFP values extracted using Metaverage, a custom-written MATLAB software. YFP/CFP values 30s before the first 21% O_2_ stimulus (baseline) were subtracted from YFP/CFP values 30s before the end of the first 21% O_2_ stimulus (stimulus). This ratio was normalised by dividing for the baseline. Values calculated for each biological replicate were entered into Prism for statistical analysis. For ADL pheromone responses average GCaMP3 intensity was calculated 5 seconds before the C9 stimulus was removed (F) and 5 seconds before the C9 stimulus was presented (F_0_). ADL pheromone-evoked Ca^2+^ responses were calculated as 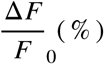. For ADL O_2_-evoked responses, a pseudo-ratio of GCaMP6s over mKate2 signal was computed to account for changes in GCaMP6s intensity due to animal movement. O_2_-evoked Ca^2+^ responses were calculated using the average GCaMP6s/mKate2 signal for a 15 sec window centred around the peak of the response (R) and a 30 sec window before stimulation with 21% O_2_ (R_0_) and expressed as 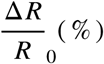.

### Cell isolation and FACS

#### Neuron isolation

Synchronised transgenic young adults in which the ADL neurons were specifically labelled using *srh-220p:mKate2* were washed 5x in M9 Buffer to remove bacteria and then dissociated as described (Beets et al., 2020; Kaletsky et al., 2018). Briefly, animals were incubated for 6.5 min in Lysis Buffer (200mM DTT, 0.25% SDS, 20mM HEPES, 3% sucrose) washed 5x in M9 Buffer, resuspended in 500 μL of 20 mg/uL Pronase in water, and pipetted up and down for 12 min at room temperature. The reaction was stopped by adding 250 μl of 2% FBS in PBS. Cells were filtered through a 5 μm syringe filter to remove clumps. mKate positive cells were sorted using a Synergy High Speed Cell Sorter (Sony Biotechnology) with gates set using a negative control prepared in parallel from dissociated unlabelled N2 animals. Positive cells were collected into 10 μL of Triton X-100 0.2% (vol/vol) supplemented with 2U/mL RNase inhibitors. Between 700 and 3000 cells were collected for each biological replicate.

### Microscopy

#### ADL neurosecretion assay

We quantified neurosecretion in young adults (< 24 hours old) expressing DAF-28::mCherry specifically in ADL neurons (*srh-220p:daf-28::mCherry*) together with a coelomocyte marker (*unc-122p*:*gfp*). To assay the effects of O_2_ levels on ADL neurosecretion we grew animals form egg to young adult either at ambient O_2_ (21% O_2_) or at 7% O_2_ using a hypoxic chamber (O_2_ Control InVitro Glove Box, Coy Laboratories). To quantify neurosecretion, we imaged the anterior pair of coelomocytes on a TE-2000 (Nikon) or a Ti2 (Nikon) wide-field microscope using a 10x air lens. Z-stack images were taken at sub-saturating exposure for both GFP and mCherry intensities and analysed using ImageJ. Coelomocytes were delineated using the GFP signal and the mCherry signal measured in the same area. Values were plotted as arbitrary units of intensity.

#### srh-220p validation

To validate the *srh-220* promoter construct we imaged wild type and *qui-1* mutants carrying a transgene expressing mKate under the control of the *srh-220* promoter (*srh-220p:mKate*). We imaged ADL cell body using a Ti2 (Nikon) wide-field microscope using a 40X air lens. Z-stack were taken avoiding saturating mKate signal. The boundary of ADL cell body was taken and intensities extracted using ImageJ. Data were plotted using Prism.

#### Dense-core vesicle localisation

To image DCVs the coding sequence of IDA-1, a DCV marker, was fused to GFP and expressed in the ADL pair of neurons using the ADL-specific promoter *srh-220*p (*srh-220p*:*ida-1::gfp*). Simultaneously, we specifically highlighted ADL by expressing cytosolic mKate (*srh-220p:mKate2*). Young adult double transgenic animals were imaged on a Ti2 (Nikon) wide-field microscope using a 40x air lens. Z-stack images were taken at sub-saturating exposure for both GFP and mKate. We delineated the boundaries for the cell body and axon of ADL using the mKate signal, and measured signals in mKate+ pixels in the GFP channel using Image J. Values were plotted in Prism, in arbitrary units.

#### QUI-1 expression and localisation

To examine the expression and sub-cellular localisation of *qui-1* we knocked in DNA encoding mNeonGreen in frame just upstream of the *qui-1* initiation codon. We imaged young adult hermaphrodites using a Ti2 (Nikon) microscope equipped with a DragonFly (Andor) spinning disk module and an EMCCD camera (iXon, Andor) with 40x or 60x objectives. Z-stacks of images acquired with sub-saturating exposure times were analysed using ImageJ.

### Analysis

#### Whole-genome sequencing

Whole-genome sequence data was analysed using a custom Python script, Cross_filter (https://github.com/lmb-seq/cross_filter). Briefly, reads were checked for quality and aligned to the *C. elegans* reference genome. Lists of mutations for each sequenced strain were then cross-referenced with a compiled list of background mutations to generate a list of strain-specific mutations.

#### RNA-sequencing

RNAseq data quality was checked using FastQC 0.11.7, before and after adaptor clipping; trimming quality was controlled using trimmomatic 0.38. Cleaned data were used for gene quantification using Salmon 1.1.0, *C. elegans* transcriptome (EnsemblMetazoa: release 46) and *C. elegans* genome (WBcel235) as decoy. We performed differential gene expression analysis using tximport 1.14.2 and DEseq2 1.26.0. Output from these programs was imported into a custom-made R program (PEAT, https://github.com/lmb-seq/PEAT) to visualise differentially expressed genes across genotypes. EnrichmentBrowser 2.16.1 was used to aggregate the enrichment of gene ontology (GO) terms from the following algorithms: Overrepresentation Analysis (ORA), Gene Set Enrichment Analysis (GSEA) and Gene Set Analysis (GSA). Beside the classic GO term annotation, we functionally annotated *C. elegans* neural genes using annotations from previously published reviews (Hobert, 2006; Robertson and Thomas, 2006) and used these in the same way as described for GO term analysis. These annotations were also used to extract data for particular classes of genes such as “Synaptic release machinery” (Figure 5A), “Chemoreceptors” (Figure 5E) and “Neuropeptide Receptors” (Figure 5B). For all Supplementary files and further analysis of RNAseq data (See Figures 5 and Figure S5) we applied an arbitrary cut off of 10 transcripts per million and q-value <0.05.

#### Tissue enrichment analysis, Heat maps and Volcano Plots

Enrichment analyses were performed using the web-based software Enrichment Analysis (https://www.wormbase.org/tools/enrichment/tea/tea.cgi) (Angeles-Albores et al., 2016). Heat maps and Volcano plots showing altered gene expression in *qui-1* mutant were generated using Prism 7 from data extracted from our custom-RNAseq analysis pipeline.

#### Statistics

Statistical tests were performed using Prism. In bar graphs error bars represent standard error of the mean (SEM). When speed plots or Ca^2+^ imaging traces are shown, shaded outlines represent the SEM. Statistics of each experiment is shown in Figure legends.

## Supplementary files

Supplementary file 1: Genes expressed in ADL neurons

Supplementary file 2: Chemoreceptors expressed in ADL neurons

Supplementary file 3: Neuropeptide receptors expressed in ADL neurons

Supplementary file 4: Neuropeptides expressed in ADL neurons

Supplementary file 5: Genes differentially regulated in ADL neurons (wild type vs *qui-1*)

Supplementary file 6: Strain list

## Acknowledgments

We would like to thank Gemma Chandratillake and Merav Cohen for identifying mutants and José David Moñino Sánchez for his help on neurosecretion assays. We are grateful to Kaveh Ashrafi (UCSF), Piali Sengupta (Brandeis) and the *Caenorhabditis* Genetic Center (funded by National Institutes of Health Infrastructure Program P40 OD010440) for strains and reagents. We thank Tim Stevens, Paula Freire-Pritchett, Alastair Crisp, Gurpreet Ghattaoraya and Fabian Amman for help with bioinformatic analysis, Ekaterina Lashmanova for help with injections, Iris Hardege for strains and Isabel Beets (KU Leuven) and members of the de Bono Lab for comments on the manuscript. We thank the CRUK Cambridge Research Institute Genomics Core for next generation sequencing and the Flow Cytometry Facility at LMB for FACS. This research was supported by the Scientific Service Units (SSU) of IST Austria through resources provided by the Bioimaging Facility (BIF), the Life Science Facility (LSF) and Scientific Computing (SciComp - Bioinformatics). This work was supported by the Medical Research Council UK (Studentship to G.V), an Advanced ERC grant (269058 ACMO to M.d.B) and a Wellcome Investigator Award (209504/Z/17/Z to M.d.B.).

## Author Contributions

G.V: Conceptualization, Investigation, Supervision, Writing – original draft, review and editing. M.d.B: Conceptualization, Investigation, Supervision, Funding acquisition, Writing – review and editing.

---

**Figure 1 – figure supplement 1.**
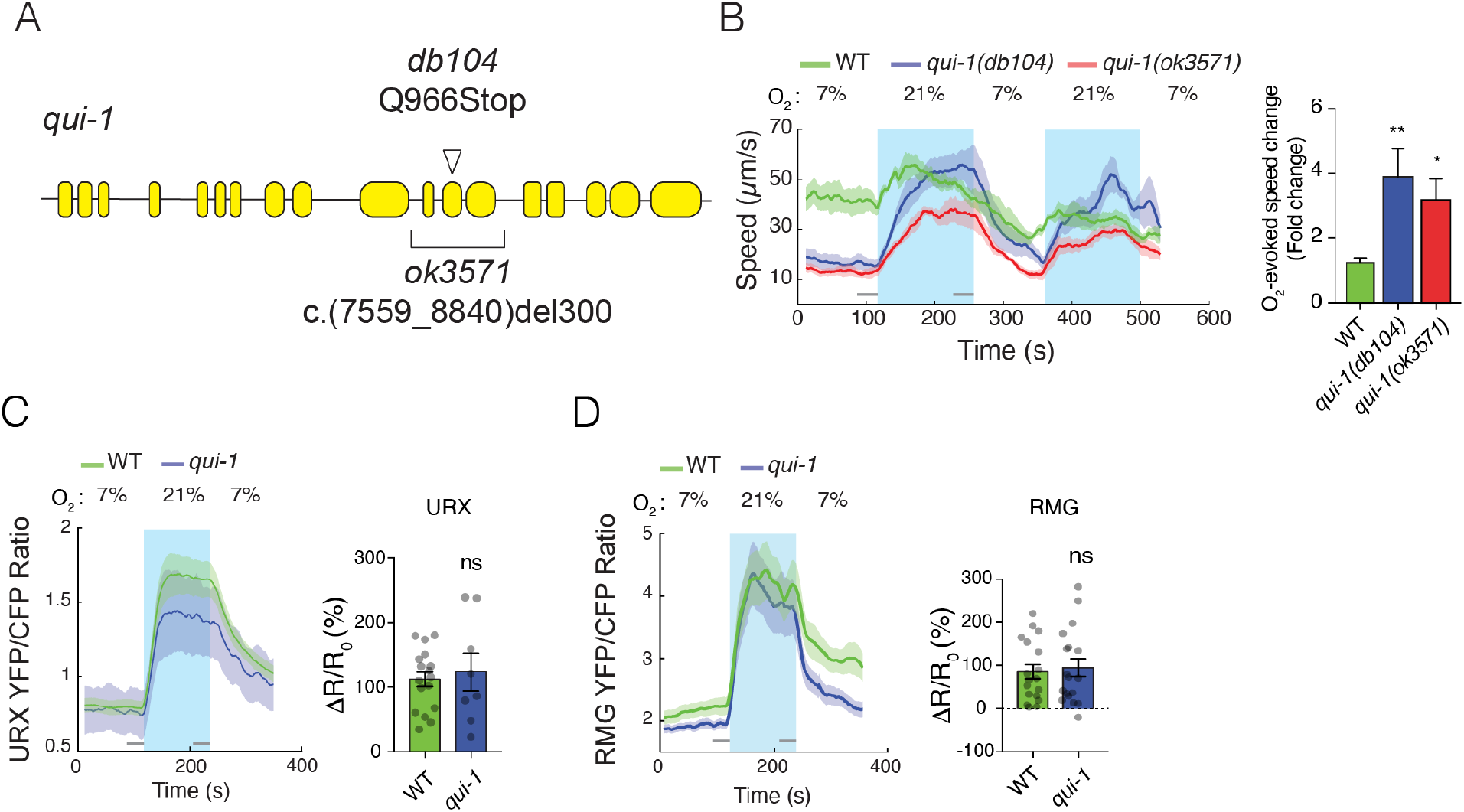
Characterisation of *qui-1* mutants. (**A**) Schematic of the *qui-1* locus, showing lesions in *qui-1*(*db104*) and *qui-1(ok3571)*. (**B**) O_2_-escape behaviors of *qui-1(db104)* and *qui-1(ok3571)* animals are indistinguishable. Left: Line shows average speed, while shading represents SEM. Grey bars show 30s time intervals used to average the animal’s speed. Right: Bar graph shows fold change in average speed at 21% O_2_ compared to 7% O_2_. N=3–5 assays. (**C-D**) O_2_-evoked Ca^2+^ responses in URX and RMG neurons are indistinguishable in *qui-1* and WT. Lines show average YFP/CFP ratio, which reports Ca^2+^ level; shading represents SEM. Bar plots quantify O_2_-evoked Ca^2+^ responses. URX N=17 (WT) and N=8 (*qui-1*), RMG N= 18 (WT) and N=18 (*qui-1*). ΔR/R_o_(%) was calculated using 30s intervals indicated by grey bars. Statistics: *, p ≤0.05, **, p ≤0.01, ***, p ≤0.001, ns=not significant, Mann-Whitney U test. Comparisons are with WT.

**Figure 2 – figure supplement 1.**
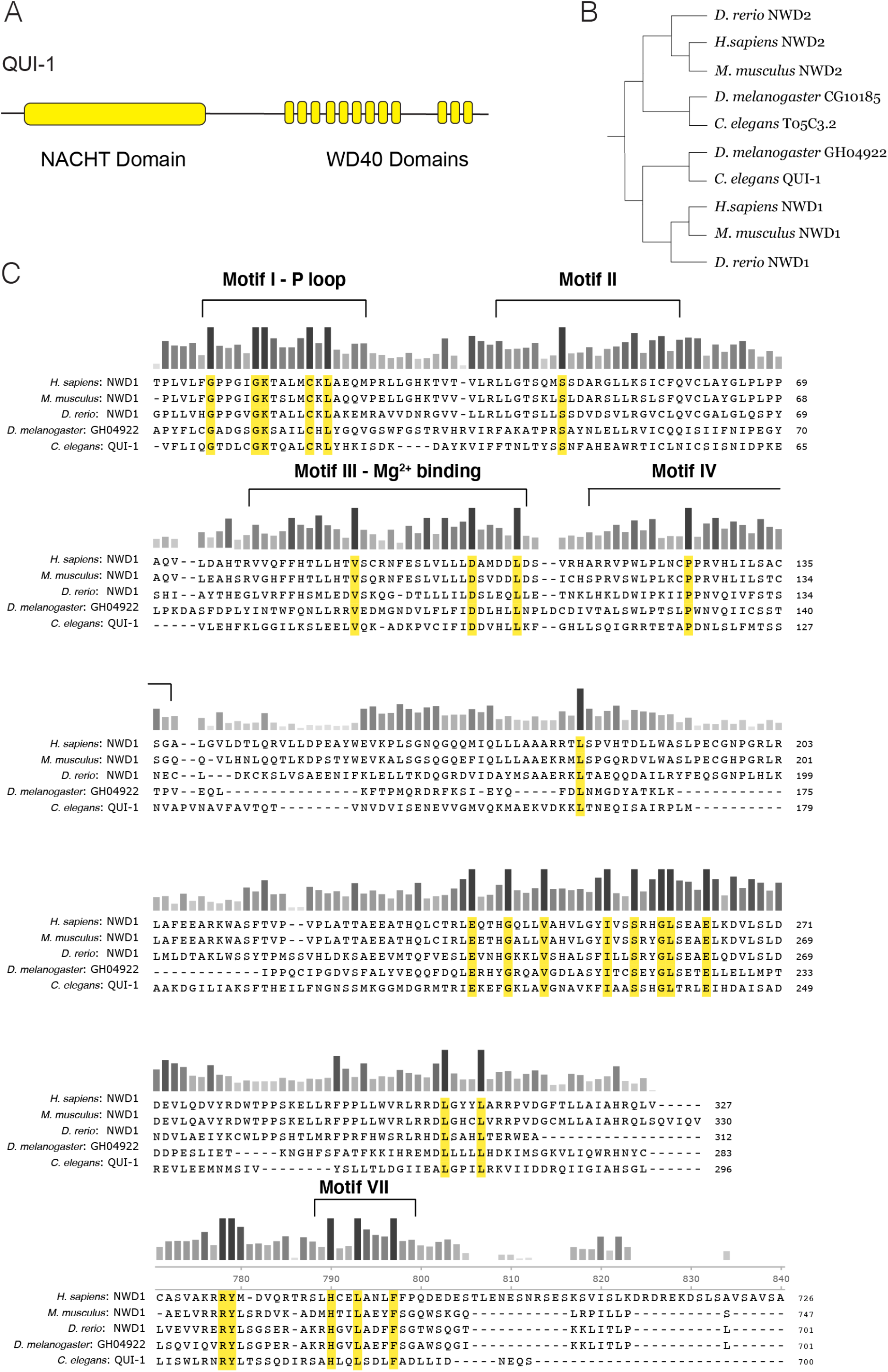
Conservation of NACHT/WD40 containing proteins. (**A**) Schematic of QUI-1 protein domains. (**B**) Phylogenetic analysis using cladogram of NWD1 and NWD2. (**C**) Sequence conservation of the QUI-1/NWD1 NACHT domain across phylogeny. Bars indicate level of conservation across species. Residues conserved in all five species are highlighted in yellow.

**Figure 3 – figure supplement 1.**
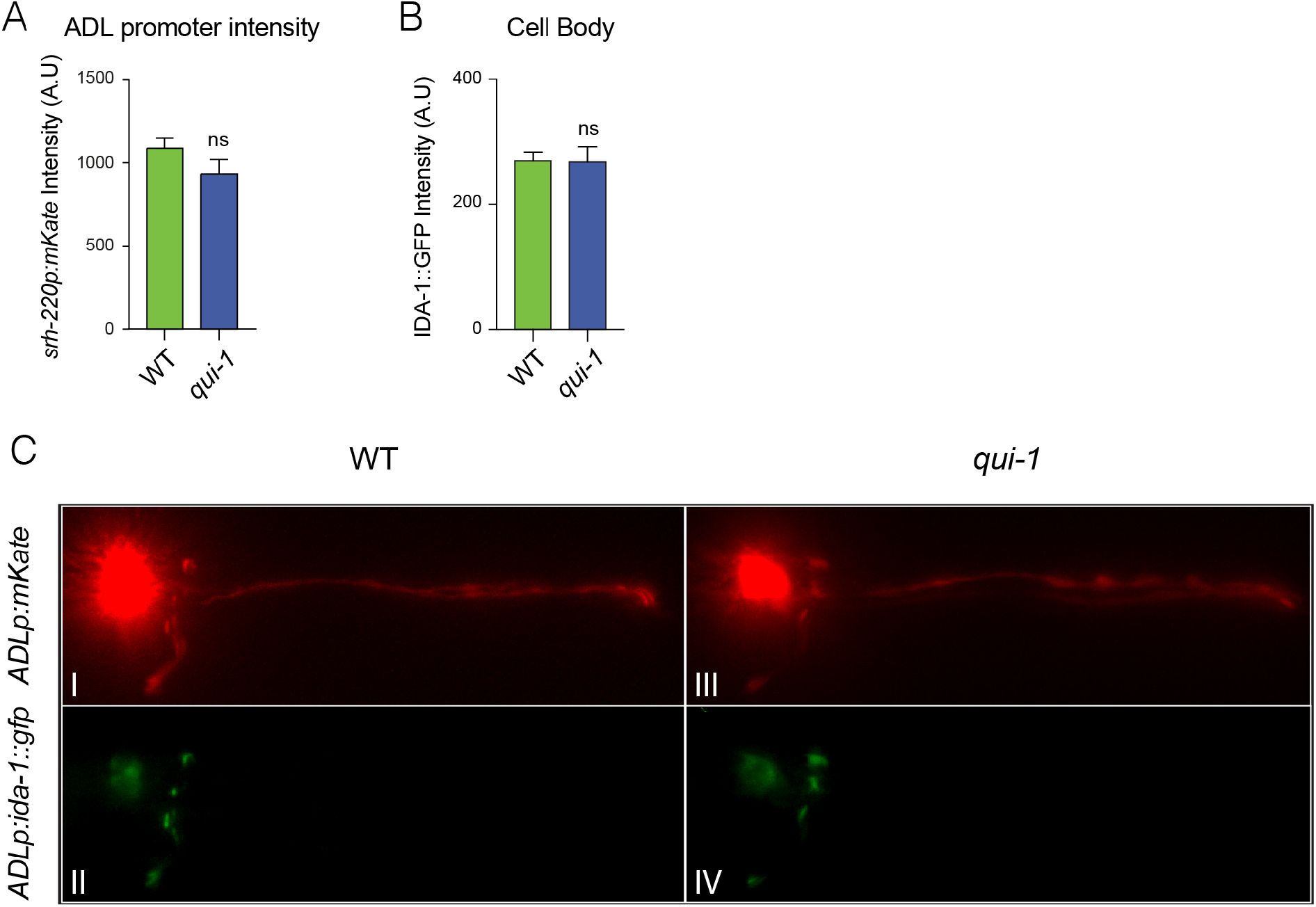
Enhanced secretion of DAF-28::mCherry from ADL reflects increased DCV release. (**A**) Expression from the ADL-specific promoter *srh-220p* is not altered in *qui-1* mutant. Bar graph shows mKate2 intensity at ADL cell body. N=27 (WT), N=21 (*qui-1*). (**B**) *qui-1* does not alter levels of the DCV marker IDA-1::GFP in ADL cell body. IDA-1::GFP was expressed in ADL (*srh-220*p) and its intensity at the cell body recorded. N=41 (WT), N=33 (*qui-1*). (**C**) IDA-1::GFP localisation in ADL in wild type and *qui-1* mutant suggests *qui-1* does not regulate overall DCVs distribution. Images are representative of images used for measurements reported in Figure 3B and Figure S3B. Statistics: ns=not significant, Mann-Whitney U test. Comparisons are with WT.

**Figure 4 – figure supplement 1.**
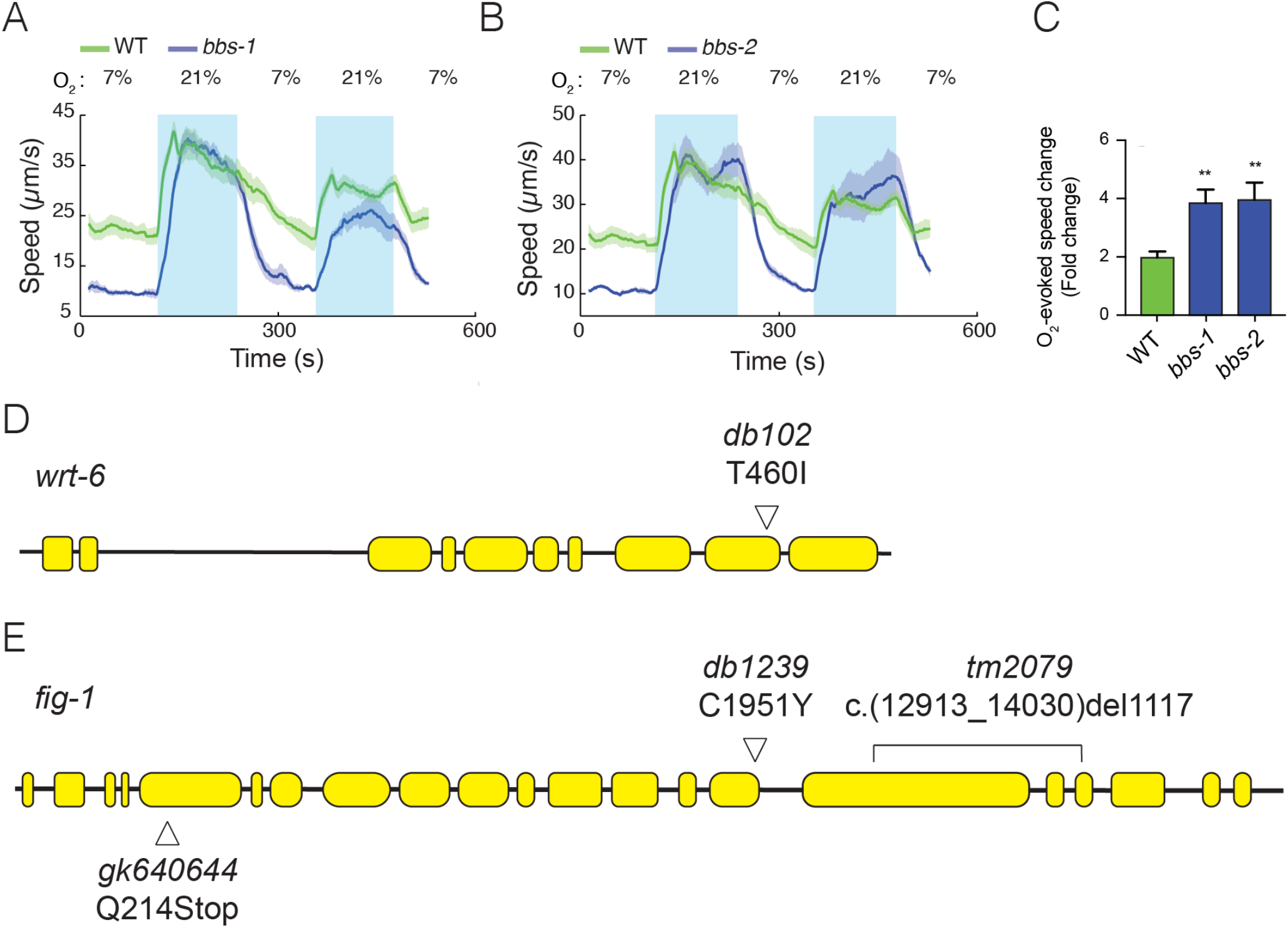
Mutants defective in subunits of the BBSome complex show enhanced O_2_-responses. (**A-C**) Mutations in two additional components of the BBSome complex, *bbs-1* (**A**), *bbs-2* (**B**), show increased O_2_-escape behavior. Left: Lines show average speed, shading represent SEM and grey bars represent 30s intervals used to average the animal’s speed. Right: Bar graph shows fold change in average speed at 21% O_2_ compared to 7% O_2_. N=5-9 assays. (**D**) Schematic of *wrt-6* locus showing the *db102* mutation isolated in our screen. The T460I substitution changes a conserved residue essential for autocleavage and activation of Hedgehog-like proteins. (**E**) Schematic of *fig-1* gene showing mutations used in Figure 4C. *fig-1*(*db102*) was isolated in our screen; *tm2079* and *gk640644* are previously isolated null alleles. Statistics: **, p value ≤0.01, Mann-Whitney U test. Comparisons are with WT.

**Figure 5 – figure supplement 1.**
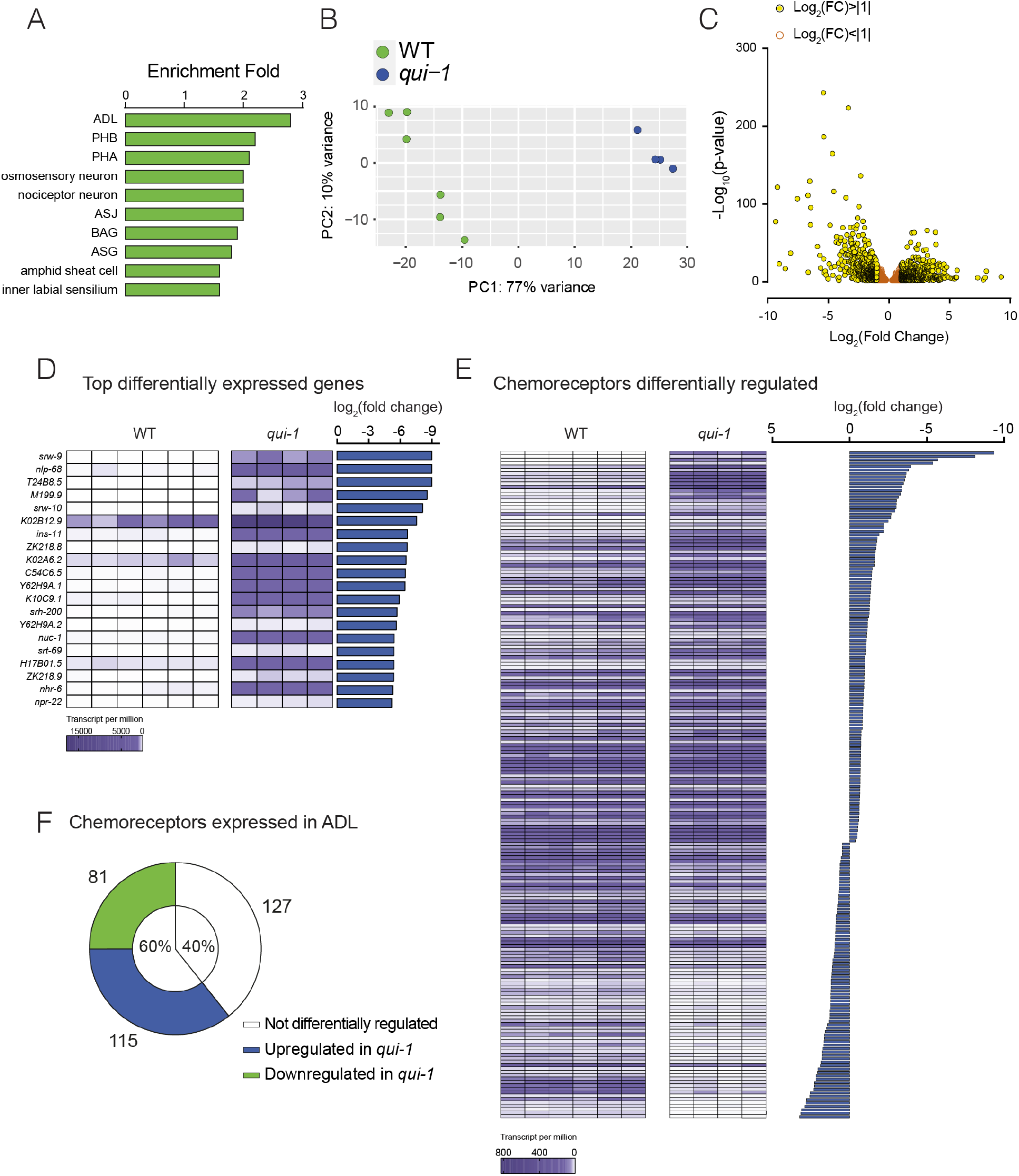
Profiling of ADL sensory neurons. **(A)** Enrichment analysis confirms our dataset most closely match the ADL profile. A list of genes expressed in ADL according to our RNAseq data (See Supplementary File 1) was analysed using the Enrichment Analysis tool (Angeles-Albores et al., 2016). **(B)** Principal component analysis showing that RNAseq data from wild type and *qui-1* biological replicates cluster by genotype. Each dot represents a single biological replicate, N=6 (WT), N=4 (*qui-1*). (**C**) Differentially expressed genes are typically up-regulated in *qui-1*. Volcano plot showing distribution of genes expressed in WT and *qui-1* according to their log_2_ fold change (x axis), and statistical significance (y axis, -log_10_(p-value)). Empty dots show genes not strongly regulated by *qui-1* (log_2_fold change between +1 and -1); yellow dots show genes strongly regulated by *qui-1* (log_2_fold change >+1 or < -1). (**D**) Heat map displays 20 most up-regulated genes in *qui-1* mutants. (**E**) Absence of *qui-1* reprograms the chemoreceptor repertoire of ADL neurons. (**D-E**) Heat maps show tpm for each gene across biological replicates. To target analysis for (E) we used lists generated in a review of the neural genome (Robertson and Thomas, 2006), and selected genes annotated as “Chemoreceptors”. All genes matching these criteria were included unless their expression was below 10 tpm in both genotypes or the q-value was >0.05 (See Supplementary File 5). (**F**) Pie chart showing absolute numbers and percentage of chemoreceptors expressed in ADL in both wild type and *qui-1* samples.

## References

Angeles-Albores D, Lee RYN, Chan J, Sternberg PW. 2016. Tissue enrichment analysis for C. elegans genomics. BMC bioinformatics 17:366–10. doi:10.1186/s12859-016-1229-9

Bacaj T, Tevlin M, Lu Y, Shaham S. 2008. Glia Are Essential for Sensory Organ Function in C. elegans. Science 322:744–747. doi:10.1126/science.1163074

Beets I, Zhang G, Fenk LA, Chen C, Nelson GM, Félix M-A, de Bono M. 2020. Natural Variation in a Dendritic Scaffold Protein Remodels Experience-Dependent Plasticity by Altering Neuropeptide Expression. Neuron 105:106-121.e10. doi:10.1016/j.neuron.2019.10.001

de Bono M, Bargmann CI. 1998. Natural variation in a neuropeptide Y receptor homolog modifies social behavior and food response in C. elegans. Cell 94:679–689.

de Bono M, Tobin DM, Davis MW, Avery L, Bargmann CI. 2002. Social feeding in Caenorhabditis elegans is induced by neurons that detect aversive stimuli. Nature 419:899–903. doi:10.1038/nature01169

Brenner S. 1974. The Genetics of Caenorhabditis elegans. Genetics 77:71–94.

Büchel C, Price C, Frackowiak RS, Friston K. 1998. Different activation patterns in the visual cortex of late and congenitally blind subjects. Brain 121:409–419. doi:10.1093/brain/121.3.409

Busch KE, Laurent P, Soltesz Z, Murphy RJ, Faivre O, Hedwig B, Thomas M, Smith HL, de Bono M. 2012. Tonic signaling from O2 sensors sets neural circuit activity and behavioral state. Nature Neuroscience 15:581–591. doi:10.1038/nn.3061

Chew YL, Tanizawa Y, Cho Y, Zhao B, Yu AJ, Ardiel EL, Rabinowitch I, Bai J, Rankin CH, Lu H, Beets I, Schafer WR. 2018. An Afferent Neuropeptide System Transmits Mechanosensory Signals Triggering Sensitization and Arousal in C. elegans. Neuron 99:1233-1246.e6. doi:10.1016/j.neuron.2018.08.003

Chronis N, Zimmer M, Bargmann CI. 2007. Microfluidics for in vivo imaging of neuronal and behavioral activity in Caenorhabditis elegans. Nature Methods 4:727–731. doi:10.1038/nmeth1075

Cook SJ, Jarrell TA, Brittin CA, Wang Y, Bloniarz AE, Yakovlev MA, Nguyen KCQ, Tang LTH, Bayer EA, Duerr JS, Bülow HE, Hobert O, Hall DH, Emmons SW. 2019. Whole-animal connectomes of both Caenorhabditis elegans sexes. Nature 571:63–71. doi:10.1038/s41586-019-1352-7

Costa WS, Yu S, Liewald JF, Gottschalk A. 2017. Fast cAMP Modulation of Neurotransmission via Neuropeptide Signals and Vesicle Loading. Curr Biol 27:495–507. doi:10.1016/j.cub.2016.12.055

Dietrich S, Hertrich I, Ackermann H. 2013. Training of ultra-fast speech comprehension induces functional reorganization of the central-visual system in late-blind humans. Front Hum Neurosci 7:701. doi:10.3389/fnhum.2013.00701

Feng G, Yi P, Yang Y, Chai Y, Tian D, Zhu Z, Liu J, Zhou F, Cheng Z, Wang X, Li W, Ou G. 2013. Developmental stage-dependent transcriptional regulatory pathways control neuroblast lineage progression. Development 140:3838–3847. doi:10.1242/dev.098723

Fenk LA, de Bono M. 2017. Memory of recent oxygen experience switches pheromone valence in Caenorhabditis elegans. Proc National Acad Sci 114:4195–4200. doi:10.1073/pnas.1618934114

Fine I, Park J-M. 2016. Blindness and Human Brain Plasticity. Annu Rev Vis Sc 4:1–20. doi:10.1146/annurev-vision-102016-061241

Gruner M, Nelson D, Winbush A, Hintz R, Ryu L, Chung SH, Kim K, Gabel CV, Linden AM van der. 2014. Feeding State, Insulin and NPR-1 Modulate Chemoreceptor Gene Expression via Integration of Sensory and Circuit Inputs. PLOS Genetics 10:e1004707. doi:10.1371/journal.pgen.1004707

Hao L, Johnsen R, Lauter G, Baillie D, Bürglin TR. 2006. Comprehensive analysis of gene expression patterns of hedgehog-related genes. BMC Genomics 7:280. doi:10.1186/1471-2164-7-280

Hilliard MA, Apicella AJ, Kerr R, Suzuki H, Bazzicalupo P, Schafer WR. 2005. In vivo imaging of C. elegans ASH neurons: cellular response and adaptation to chemical repellents. The EMBO Journal 24:63–72. doi:10.1038/sj.emboj.7600493

Hilliard MA, Bergamasco C, Arbucci S, Plasterk RH, Bazzicalupo P. 2004. Worms taste bitter: ASH neurons, QUI-1, GPA-3 and ODR-3 mediate quinine avoidance in Caenorhabditis elegans. The EMBO Journal 23:1101–1111. doi:10.1038/sj.emboj.7600107

Hobert O. 2006. The neuronal genome of Caenorhabditis elegans.Undefined, WormBook?: The Online Review of C. Elegans Biology. pp. 1–106. doi:10.1895/wormbook.1.161.1

Jang H, Kim K, Neal SJ, Macosko E, Kim D, Butcher RA, Zeiger DM, Bargmann CI, Sengupta P. 2012. Neuromodulatory State and Sex Specify Alternative Behaviors through Antagonistic Synaptic Pathways in C. elegans. Neuron 75:585–592. doi:10.1016/j.neuron.2012.06.034

Jarrell TA, Wang Y, Bloniarz AE, Brittin CA, Xu M, Thomson JN, Albertson DG, Hall DH, Emmons SW. 2012. The Connectome of a Decision-Making Neural Network. Science 337:437–444. doi:10.1126/science.1221762

Kaletsky R, Yao V, Williams A, Runnels AM, Tadych A, Zhou S, Troyanskaya OG, Murphy CT. 2018. Transcriptome analysis of adult Caenorhabditis elegans cells reveals tissue-specific gene and isoform expression. PLOS Genetics 14:e1007559–29. doi:10.1371/journal.pgen.1007559

Laurent P, Soltesz Z, Nelson GM, Chen C, Arellano-Carbajal F, Levy E, de Bono M. 2015. Decoding a neural circuit controlling global animal state in C. elegans. eLife 4:e1004156. doi:10.7554/elife.04241

Lee BH, Ashrafi K. 2008. A TRPV Channel Modulates C. elegans Neurosecretion, Larval Starvation Survival, and Adult Lifespan. PLOS Genetics 4:e1000213. doi:10.1371/journal.pgen.1000213

Lee BH, Liu J, Wong D, Srinivasan S, Ashrafi K. 2011. Hyperactive Neuroendocrine Secretion Causes Size, Feeding, and Metabolic Defects of C. elegans Bardet-Biedl Syndrome Mutants. PLOS Biol 9:e1001219. doi:10.1371/journal.pbio.1001219

Lee H-K, Whitt JL. 2015. Cross-modal synaptic plasticity in adult primary sensory cortices. Curr Opin Neurobiol 35:119–126. doi:10.1016/j.conb.2015.08.002

Liu P, Lechtreck KF. 2018. The Bardet–Biedl syndrome protein complex is an adapter expanding the cargo range of intraflagellar transport trains for ciliary export. Proceedings of the National Academy of Sciences of the United States of America 115:E934–E943. doi:10.1073/pnas.1713226115

Macosko EZ, Pokala N, Feinberg EH, Chalasani SH, Butcher RA, Clardy J, Bargmann CI. 2009. A hub-and-spoke circuit drives pheromone attraction and social behaviour in C. elegans. Nature 458:1171–1175. doi:10.1038/nature07886

Merabet LB, Pascual-Leone A. 2009. Neural reorganization following sensory loss: the opportunity of change. Nature Reviews Neuroscience 11:44–52. doi:10.1038/nrn2758

Mok CA, Healey MP, Shekhar T, Leroux MR, Héon E, Zhen M. 2011. Mutations in a Guanylate Cyclase GCY-35/GCY-36 Modify Bardet-Biedl Syndrome–Associated Phenotypes in Caenorhabditis elegans. PLOS Genetics 7:e1002335. doi:10.1371/journal.pgen.1002335

Moreno E, Sieriebriennikov B, Witte H, delsperger CR x000F6, Lightfoot JW, Sommer RJ. 2017. Regulation of hyperoxia-induced social behaviour in Pristionchus pacificus nematodes requires a novel cilia-mediated environmental input. Scientific Reports 1–13. doi:10.1038/s41598-017-18019-0

Moreno E, Sommer RJ. 2018. A cilia-mediated environmental input induces solitary behaviour in Caenorhabditis elegans and Pristionchus pacificus nematodes. Nematology 20:201–209. doi:10.1163/15685411-00003159

Neal SJ, Park J, DiTirro D, Yoon J, Shibuya M, Choi W, Schroeder FC, Butcher RA, Kim K, Sengupta P. 2016. A Forward Genetic Screen for Molecules Involved in Pheromone-Induced Dauer Formation in Caenorhabditis elegans. G3 (Bethesda, Md) 6:1475–1487. doi:10.1534/g3.115.026450

Petrus E, Isaiah A, Jones AP, Li D, Wang H, Lee H-K, Kanold PO. 2014. Crossmodal Induction of Thalamocortical Potentiation Leads to Enhanced Information Processing in the Auditory Cortex. Neuron 81:664–673. doi:10.1016/j.neuron.2013.11.023

Pocock R, Hobert O. 2010. Hypoxia activates a latent circuit for processing gustatory information in C. elegans. Nat Neurosci 13:610–614. doi:10.1038/nn.2537

Rabinowitch I, Laurent P, Zhao B, Walker D, Beets I, Schoofs L, Bai J, Schafer WR, Treinin M. 2016. Neuropeptide-Driven Cross-Modal Plasticity following Sensory Loss in Caenorhabditis elegans. PLOS Biol 14:e1002348–24. doi:10.1371/journal.pbio.1002348

Rauschecker JP. 1995. Compensatory plasticity and sensory substitution in the cerebral cortex. Trends Neurosci 18:36–43. doi:10.1016/0166-2236(95)93948-w

Robertson HM, Thomas JH. 2006. The putative chemoreceptor families of C. elegans. Wormbook 1–12. doi:10.1895/wormbook.1.66.1

Rogers C, Reale V, Kim K, Chatwin H, Li C, Evans P, de Bono M. 2003. Inhibition of Caenorhabditis elegans social feeding by FMRFamide-related peptide activation of NPR-1. Nature Neuroscience 6:1178–1185. doi:10.1038/nn1140

Sadato N, Pascual-Leone A, Grafman J, Ibañez V, Deiber M-P, Dold G, Hallett M. 1996. Activation of the primary visual cortex by Braille reading in blind subjects. Nature 380:526–528. doi:10.1038/380526a0

Saeki S, Yamamoto M, Iino Y. 2001. Plasticity of chemotaxis revealed by paired presentation of a chemoattractant and starvation in the nematode Caenorhabditis elegans. J Exp Biology 204:1757–64.

Tan PL, Barr T, Inglis PN, Mitsuma N, Huang SM, Garcia-Gonzalez MA, Bradley BA, Coforio S, Albrecht PJ, Watnick T, Germino GG, Beales PL, Caterina MJ, Leroux MR, Rice FL, Katsanis N. 2007. Loss of Bardet Biedl syndrome proteins causes defects in peripheral sensory innervation and function. Proceedings of the National Academy of Sciences of the United States of America 104:17524–17529. doi:10.1073/pnas.0706618104

Tyssowski KM, DeStefino NR, Cho J-H, Dunn CJ, Poston RG, Carty CE, Jones RD, Chang SM, Romeo P, Wurzelmann MK, Ward JM, Andermann ML, Saha RN, Dudek SM, Gray JM. 2018. Different Neuronal Activity Patterns Induce Different Gene Expression Programs. Neuron 98:530-546.e11. doi:10.1016/j.neuron.2018.04.001

Vidal B, Aghayeva U, Sun H, Wang C, Glenwinkel L, Bayer EA, Hobert O. 2018. An atlas of Caenorhabditis elegans chemoreceptor expression. PLOS Biol 16:e2004218–34. doi:10.1371/journal.pbio.2004218

Wanet-Defalque M-C, Veraart C, Volder AD, Metz R, Michel C, Dooms G, Goffinet A. 1988. High metabolic activity in the visual cortex of early blind human subjects. Brain Res 446:369–373. doi:10.1016/0006-8993(88)90896-7

White JG, Southgate E, Thomson JN, Brenner S. 1986. The Structure of the Nervous System of the Nematode Caenorhabditis elegans. Philosophical Transactions of the Royal Society of London B: Biological Sciences 314:1–340. doi:10.1098/rstb.1986.0056

Wu J, Duggan A, Chalfie M. 2001. Inhibition of touch cell fate by egl-44 and egl-46 in C. elegans. Gene Dev 15:789–802. doi:10.1101/gad.857401

Yap E-L, Greenberg ME. 2018. Activity-Regulated Transcription: Bridging the Gap between Neural Activity and Behavior. Neuron 100:330–348. doi:10.1016/j.neuron.2018.10.013

Zhang Y, Lu H, Bargmann CI. 2005. Pathogenic bacteria induce aversive olfactory learning in Caenorhabditis elegans. Nature 438:179–184. doi:10.1038/nature04216

Zimmer M, Gray JM, Pokala N, Chang AJ, Karow DS, Marletta MA, Hudson ML, Morton DB, Chronis N, Bargmann CI. 2009. Neurons detect increases and decreases in oxygen levels using distinct guanylate cyclases. Neuron 61:865–879. doi:10.1016/j.neuron.2009.02.013

Zuryn S, Gras SL, Jamet K, Jarriault S. 2010. A Strategy for Direct Mapping and Identification of Mutations by Whole-Genome Sequencing. Genetics 186:427–430. doi:10.1534/genetics.110.119230

